# High-resolution mapping of cis-regulatory variation in budding yeast

**DOI:** 10.1101/143164

**Authors:** Ryosuke Kita, Sandeep Venkataram, Yiqi Zhou, Hunter B. Fraser

## Abstract

Genetic variants affecting gene expression levels are a major source of phenotypic variation. The approximate locations of these variants can be mapped as expression quantitative trait loci (eQTLs); however a major limitation of eQTLs is their low resolution, which precludes investigation of the causal variants and their molecular mechanisms. Here we report RNA-seq and full genome sequences for 85 diverse isolates of the yeast *Saccharomyces cerevisiae*—including wild, domesticated, and human clinical strains—which allowed us to perform eQTL mapping with 50-fold higher resolution than previously possible. In addition to variants in promoters, we uncovered an important role for variants in 3’ untranslated regions, especially those affecting binding of the PUF family of RNA-binding proteins. The eQTLs are predominantly under purifying selection, particularly those affecting essential genes and conserved genes. However, applying the sign test for lineage-specific selection revealed the polygenic up-regulation of dozens of biofilm suppressor genes in strains isolated from human patients, consistent with the key role of biofilms in fungal pathogenicity. In addition, a single variant in the promoter of a biofilm suppressor, *NIT3,* showed the strongest genome-wide association with clinical origin. Altogether our results demonstrate the power of high-resolution eQTL mapping in understanding the molecular mechanisms of regulatory variation, as well as the natural selection acting on this variation that drives adaptation to environments ranging from laboratories to vineyards to the human body.

## INTRODUCTION

Genome-wide association studies (GWAS) have identified thousands of associations between genetic variants and phenotypes in a wide range of species. As more of these associations are identified, there is a concomitantly increasing need to answer the question of *how* these variants shape particular phenotypes. A common mechanism for these phenotype-altering variants is via changes in gene expression (Musunuru et al. 2010; Albert and Kruglyak 2015; Claussnitzer et al. 2015). Indeed, human disease-associated variants are highly enriched for regulatory functions, and genetic variants associated with gene expression can implicate causal genes (Schaub et al. 2012; Gusev et al. 2016; Zhu et al. 2016). In addition, gene expression regulation has been found to be the predominant target of positive selection in recent human evolution (Fraser 2013; Enard et al. 2014). These findings all support the longstanding hypothesis that cis-regulatory variants are a critical component in the evolution of complex traits.

The budding yeast *Saccharomyces cerevisiae* is a key model organism for investigating how genetic variants influence gene expression. Genetic variants or loci associated with a gene’s mRNA abundance are known as expression quantitative trait loci, or eQTLs. Recombinant lines of two *S. cerevisiae* strains were utilized for the first genome-wide eQTL mapping (Brem et al. 2002). This work identified widespread “local” eQTLs located very close to the regulated gene, which are predominantly caused by cis-acting variants (Ronald et al. 2005), as well as several trans-acting “hotspots” where a single locus controls the expression of many genes (Brem et al. 2002). Additional studies on the same genetic cross identified gene-by-environment interactions of eQTLs (Smith and Kruglyak 2008), genetic interactions between eQTLs resulting in non-additive effects (Brem et al. 2005), and widespread adaptive evolution of gene expression (Fraser et al. 2010).

*Saccharomyces* hybrids have also proven to be a valuable resource to study *cis*-regulatory variation, since allele-specific expression (ASE) in a hybrid reflects only cis-acting divergence (Ronald et al. 2005; Tirosh et al. 2009), as opposed to *trans-acting* divergence that affects both alleles. Studies of hybrid ASE have revealed interesting cases of pathway-level regulatory divergence (Martin et al. 2012; Roop et al. 2016) and examples of polygenic adaptation affecting traits such as pathogenicity, ergosterol biosynthesis, and toxin resistance (Fraser et al. 2012; Chang et al. 2013; Naranjo et al. 2015). A key difference between hybrid ASE and eQTL mapping is that ASE reveals the genes affected by cis-regulatory variation, but does not indicate the location of the causal variant as eQTL mapping can.

Despite the utility of eQTL studies in *S. cerevisiae,* they have had several limitations. First, existing eQTLs generally span many kilobases, and thus cannot determine the precise location of the causal variant. This is because the mapping was performed with first-generation meiotic segregants with a limited number of recombinations between parental genomes; these recombination breakpoints are needed to map QTLs, since only when the linked alleles are separated by recombination can their effects be distinguished. Second, yeast eQTLs have been mapped from genetic crosses between just a few parental strains, and thus do not sample most of the natural variation across the species. Incorporating a diverse collection of strains to map eQTLs may thus allow not only greater mapping resolution, but also a deeper understanding of species-wide patterns of natural selection on regulatory variation.

Here we address both of these limitations by mapping eQTLs in a diverse set of 85 *S. cerevisiae* strains. By performing eQTL mapping in a wide variety of genetic backgrounds, we minimized linkage disequilibrium between nearby variants and thus mapped eQTLs with high resolution, similar to GWAS in humans and other species (Altshuler et al. 2009, Flint and Eskin 2012). In addition, these 85 strains were isolated from a wide range of ecological niches including clinical isolates, laboratory strains, and vineyard isolates, and thus allowed us to investigate the evolution of gene expression regulation across these diverse environments.

## RESULTS

### DNA and RNA-seq of 85 yeast isolates

We obtained 85 *S. cerevisiae* strains from across the world, which included domesticated, wild, and clinical isolates (Figure 1A, Supplemental Table 1) (Muller et al. 2011). The domesticated isolates are laboratory strains and vineyard strains; the wild isolates were obtained from wild plants, such as oak trees; and the clinical strains were isolated from sites of infection in human patients. Using these isolates, Muller *et al.* identified 135,771 polymorphisms using a microarray. Because the microarray cannot identify the precise variants due to its limited resolution, we performed whole-genome sequencing of the 85 isolates (Materials and Methods, Supplemental Figure 1). Sixty-five of these isolates were recently sequenced independently (Strope *et al.* 2015), and we observed strong concordance of variant calls for these strains (Materials and Methods, Supplemental Figure 2). Using variants within conserved chromosomal regions, we created a phylogenetic tree (Figure 1B, Supplementary Text). As previously observed, isolates with similar ecological niches exhibited a moderate degree of clustering (Liti et al. 2009; Schacherer et al. 2009; Strope et al. 2015) (Supplemental Figure 3). The population structure was also associated with geographical origin, e.g. with a tightly grouped set of vineyard isolates from Italy, but there were also clusters of similar ecological niche that spanned multiple continents (Supplemental Figure 3).

**Figure 1:**
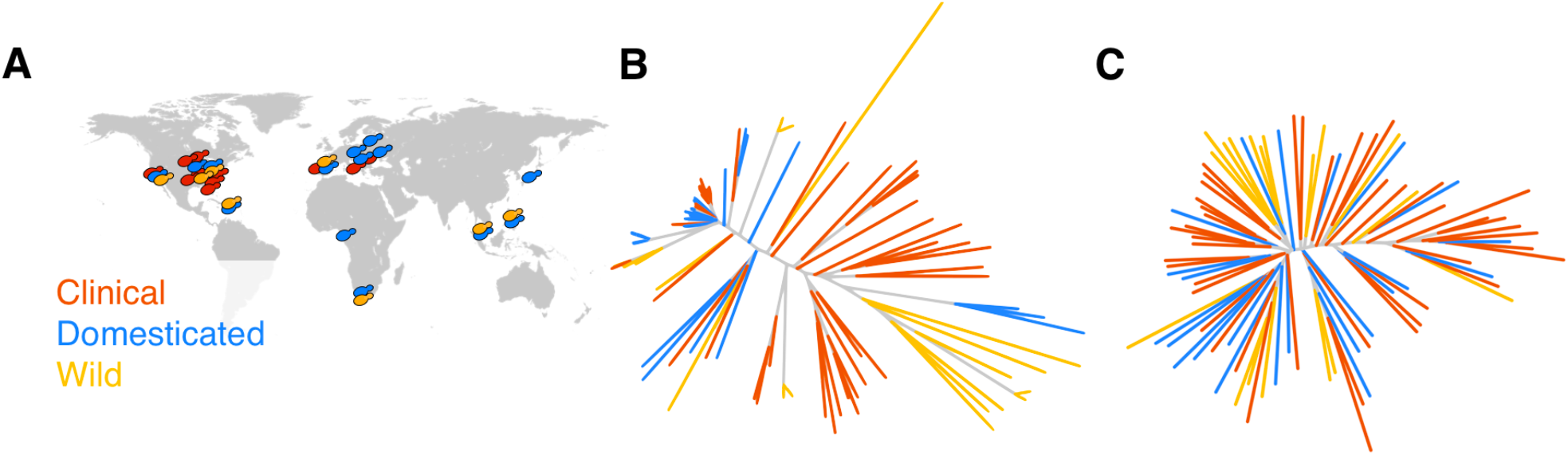
Origins and relationships among 85 isolates. (A) The isolation locations of the strains analyzed in this study. Colors represent the ecological niche of origin. (B) Neighbor-joining tree created using a 218 kb sequence comprised of conserved regions across the genome. Colors label the origin of the strain. (C) Neighbor-joining tree created using the Euclidean distance of the normalized expression across all genes.

To measure gene expression levels across all 85 isolates, we performed RNA-seq in YPD at 30° C (Materials and Methods, Supplemental Figure 4). To control for potential read mapping biases that could cause reads with non-reference alleles to map less well, we masked the reference genome prior to mapping, converting all single-nucleotide variants identified in our 85 strains to N’s. We found no association between mapping rates and divergence from the reference genome, suggesting that mapping bias was not a major issue (Supplemental Figure 4). Using both DNA- and RNA-seq data, we also assessed whether any genes from the reference genome were absent in the genome of each isolate (Supplementary Text, Supplemental Figure 5). We found a median of 30 genes that were missing across the isolates, located primarily in the sub-telomeric regions (Supplemental Figure 6). These genes were removed from eQTL mapping analyses described below.

Because we identified population structure based on ecological niche in the genomic variants, we asked whether the transcriptomes also exhibit this structure. To test this, we computed the Euclidean distance across all gene expression levels between every pair of isolates, and used these to construct a neighbor-joining tree (Figure 1C). Isolates from the same ecological niche were visibly less clustered together than in the genomic tree, with the large clinical and wild clades split up and interspersed with isolates from other niches. To quantify these relationships, we calculated the first six principal components of the gene expression profiles (Materials and Methods). We found only a weak association between the principal components and ecological origin or sequencing batch, suggesting that these are not the primary determinants of these transcriptomes (Supplemental Figure 7, Supplemental Figure 8). On the other hand, the principal components calculated from the genotypes exhibited strong associations with ecological origin (Supplemental Figure 9).

### High resolution eQTL mapping

To investigate the genetic basis of transcription across these isolates, we mapped the expression level of each gene to variants within 2.5 kb of the transcript boundaries (Materials and Methods: “eQTL mapping”). We used GEMMA (Zhou and Stephens 2012) to perform this mapping, as it has been shown to adequately control for population structure in *S. cerevisiae* local eQTL mapping (Connelly and Akey 2012). We discovered 1403 genes with a local eQTL at FDR < 0.05. To replicate these eQTLs, we compared them with a previous study that performed RNA-seq on 22 yeast isolates (Skelly et al. 2013). We performed the eQTL analysis using the same method as above, and found significant enrichment of low p-values and concordant directionality with our eQTLs (Supplemental Figure 10, Supplemental Text).

Because we mapped eQTLs using a large number of isolates with high genetic diversity, we hypothesized that our mapping resolution might be higher than previous eQTL analyses in *S. cerevisiae*. To test this, we compared the resolution of the eQTLs from this study with the resolution from a previous analysis of 112 segregants from a genetic cross (Ronald et al. 2005). QTL widths are typically reported as a 1-LOD (logarithm-of-odds) or 2-LOD support interval (defined as the distance between two genetic markers: the first marker to the left of the most significant marker in a QTL to have a LOD score at least 1 or 2 lower than the most significant marker, and the equivalent marker to the right). We observed a 49.7-fold higher resolution in our analysis with a median width of 1210 bp, compared to the median of 60,230 bp in the segregant analysis (p < 10^−15^, Figure 2A; see Materials and Methods). We replicated this finding with 2-LOD support intervals from the same study, and 1.5 LOD support intervals from another yeast eQTL study (Smith and Kruglyak 2008, Supplemental Figure 12A).

**Figure 2:**
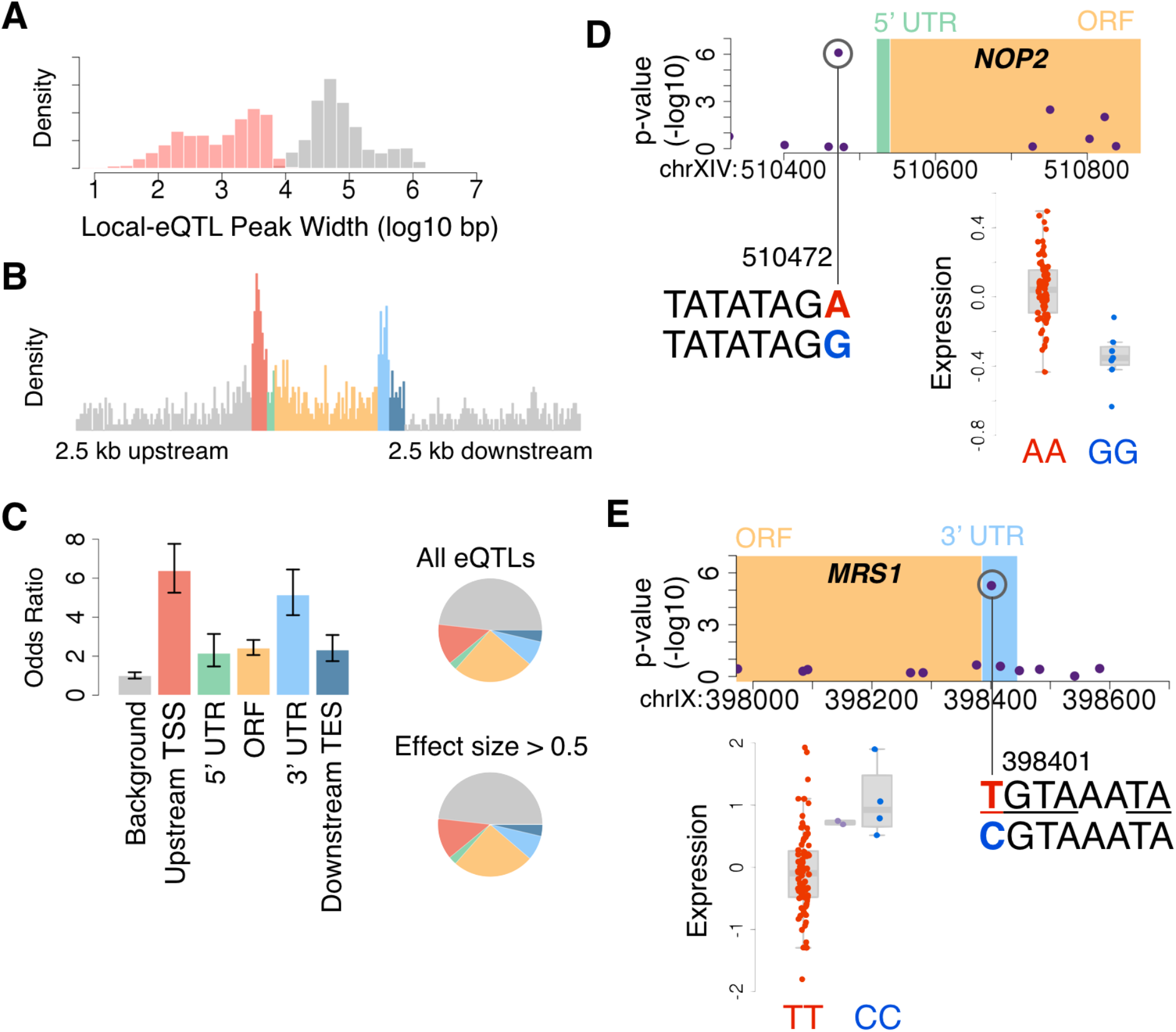
High-resolution eQTL mapping. (A) Distribution of eQTL resolution width in the segregant analysis (grey) and this study (red). (B) Distribution of eQTLs relative to transcript components labeled in panel C. The upstream TSS and downstream TES refer to 1 kb windows. The length of each component has been scaled (set to the mean length of that component across all genes) so that all genes are displayed with the same length for each component. (C) Odds ratio of eQTL density of transcript component relative to expected. Pie chart of the proportion of transcript components for all eQTLs and eQTLs with large effect size. (D, E) eQTL for *NOP2* and *MRS1.* Purple points indicate the p-value for each variant that was tested. The letters indicate the sequence of the region neighboring the variant with the most significant p-value, and the colored letters indicate the alleles for that variant. The box plot depicts the normalized expression levels for isolates with each allele.

Taking advantage of this high resolution, we investigated the location of the strongest eQTL for each gene relative to several transcript components: the transcription start site (TSS), 5’ untranslated region (UTR), open reading frame (ORF), 3’ UTR, and transcription end site. Previous analysis of 22 yeast strains did not detect enrichment of eQTLs in a particular component (Skelly et al. 2013). We found the strongest enrichments upstream of the TSS (i.e. the promoter) and within the 3’ UTR (Figure 2B). To account for SNP density differences between components, we calculated the odds ratio of the enrichment compared to that expected by chance, which produced similar results (Figure 2C). To evaluate whether variants in some locations might be more likely to result in strong effects on transcript levels, we examined the locations of eQTLs with large effect size (beta > 0.5, 176 genes, Figure 2C). We found that the locations were similar to weaker eQTLs, suggesting that large-effect eQTLs are no more likely to occur in a particular location than weak eQTLs (Supplemental Figure 12B).

An example of our high-resolution eQTL mapping is an eQTL for the gene *NOP2* (Figure 2D). The strongest associating variant for *NOP2* was 50 bp upstream of the TSS, and the neighboring SNPs had far weaker associations, allowing us to define the likely causal SNP. Notably, this variant is found in a TATA-element as determined by a ChIP-exo study of pre-initiation complex protein binding (Rhee and Pugh 2012). Another illustrative example is an eQTL for the gene *MRS1.* Here, the strongest associating variant is within the 3’ UTR, and the neighboring SNPs also have much weaker associations. This variant is located within an RNA-binding motif for PUF proteins: UGUA--UA (Hogan et al. 2015). In addition, Puf3p binds to this transcript (Gerber et al. 2004), and this variant is found 4 bases away from a Puf3p binding site identified by PAR-Clip (Freeberg et al. 2013).

Surprisingly, the 3’ UTR enrichment is almost as strong as the enrichment upstream of the TSS (Figure 2C). The 3’ UTR is the site of many mRNA-protein interactions that affect RNA decay (Pai et al. 2012), so we thus investigated whether there was any enrichment for particular binding sites among the 3’ UTR eQTLs. We performed a de novo differential motif analysis in these regions, where we controlled for motifs that are enriched in 3’ UTRs overall by using all 3’ UTRs as the background (Supplementary Text). This analysis identified two significant motifs, which contained the RNA-binding motifs of Puf3p (UGUA) and Puf2p (UAAU) (Gerber et al. 2008) (Supplemental Figure 13). Performing a similar de novo motif analysis on the promoter regions, we found strong enrichment of several motifs, although we do not yet know what functions these may have (Supplemental Figure 14).

In addition to the enrichment in the 3’ UTR and promoter regions, we also identified 310 eQTLs in the ORFs of their target genes, with increasing enrichment near the 5’ end. Regulatory variants in the ORF may have either a *cis*-acting mechanism (such as disruption of transcription factor binding), or a *trans*-acting effect (such as a non-synonymous mutation affecting a gene’s auto-regulation, as seen for *AMN1)* (Ronald et al. 2005). If such a self-regulating *trans*-mechanism was a common effect, we would expect an enrichment of non-synonymous variants, since these are more likely to disrupt protein function than synonymous variants. Among all eQTLs within ORFs, we found no enrichment of non-synonymous compared to synonymous variants (67.7% synonymous eQTL variants observed, compared to 66.9% expected by chance based on all variants in these genes), suggesting that most eQTLs within ORFs are unlikely to act in *trans*.

### Species-wide selection pressures on eQTLs

Gene expression has been shown to be predominantly under stabilizing selection in several species (Denver et al. 2005; Rifkin et al. 2005; Landry et al. 2007), though positive selection is also acting on the expression of many genes in yeast (Fraser et al. 2010). To understand the evolutionary pressures on the eQTLs in this study, we evaluated the eQTLs for overall selection using two metrics: 1) The allele frequencies of the eQTLs and 2) The presence and absence of eQTLs. Previous studies have found that eQTLs in humans and *Capsella grandiflora* are enriched for rare alleles, suggesting that eQTLs are under purifying selection (Battle et al. 2014; Josephs et al. 2015). In yeast, a similar pattern of broad purifying selection has been observed (Ronald and Akey 2007); however because only three *S. cerevisiae* genomes were available at the time, rare alleles could not be confidently identified (Ronald and Akey 2007). With many more genomes and a wider sampling of gene expression variation, we revisited this conclusion with increased power and resolution.

To identify whether eQTLs are either depleted or enriched for rare alleles, a null distribution of expected minor allele frequencies (MAFs) is essential. To account for the statistical bias that eQTLs are more easily detected at high MAF, we generated a null distribution of MAFs using the same strategy as a previous analysis (Josephs et al. 2015) (see Supplementary Text). Comparing the MAF distribution of the randomized vs. real eQTLs, we found an enrichment of eQTLs among rare alleles (MAF < 0.15, p<10^−3^, Figure 3A), and a depletion of eQTLs among common alleles (MAF > 0.45, p<10^−3^, Figure 3A). These analyses confirm that the dominant selective force on eQTLs in *S. cerevisiae* is negative selection.

**Figure 3:**
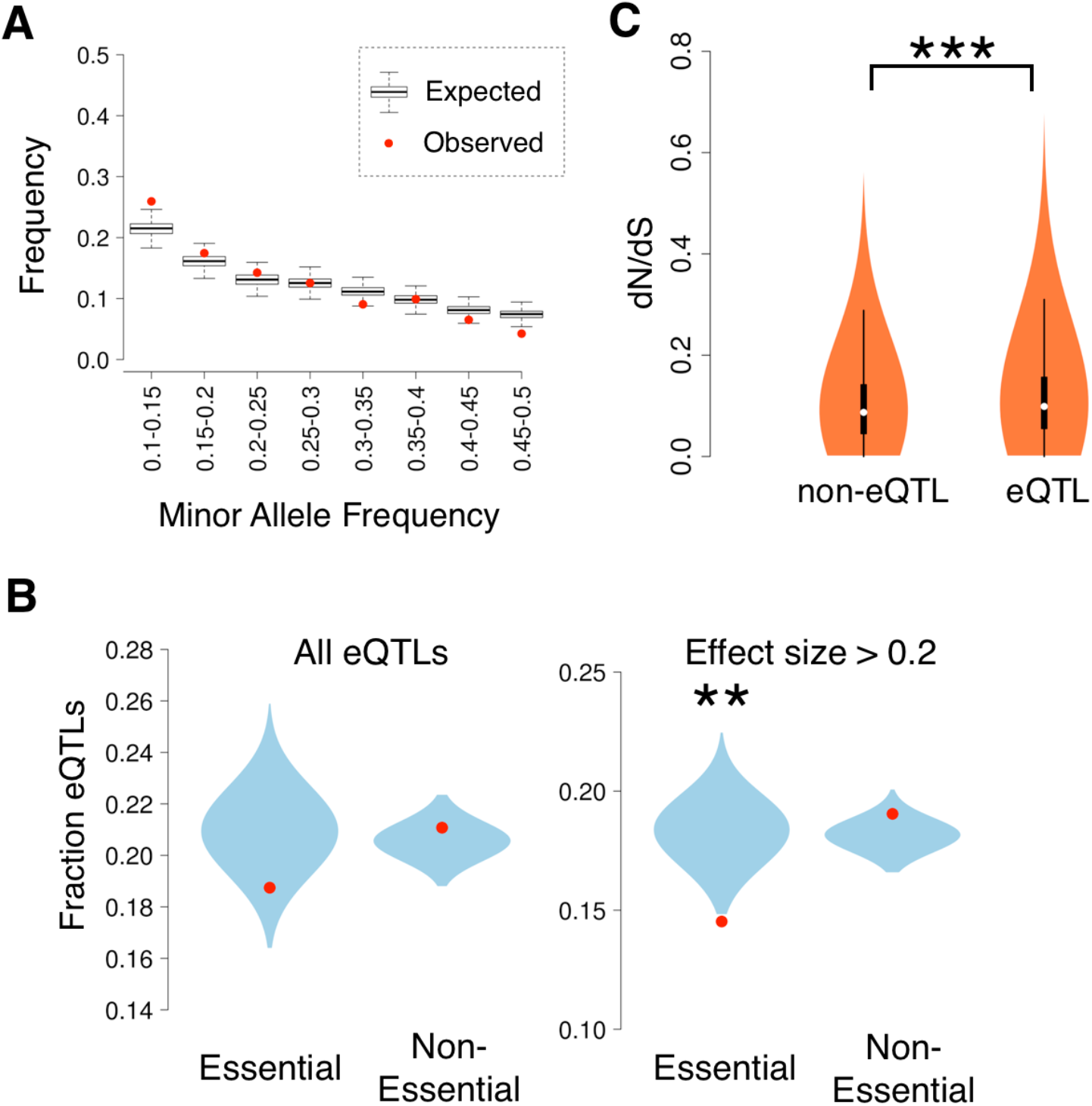
Selection on eQTLs by changes in presence/absence and allele frequency. (A) The proportion of eQTLs observed within the specified minor allele frequency range is plotted in red. The boxplots represent the null distribution of expected minor allele frequencies. (B) The observed fraction of eQTLs among the genes is plotted in red. The blue violin plots depict the distribution of expected fraction of eQTLs from permutations that control for expression level. ** indicates permutation p < 10^−3^ and * indicates p < 0.05. (C) The distribution of dN/dS values of genes as violin plots segregated by presence and absence of eQTLs. *** indicates p-value < 10^−4^.

We then hypothesized that the regulation of essential genes may be under greater purifying selection than nonessential genes (Fraser et al. 2004; Tirosh and Barkai 2008). If so, mutations affecting expression of these genes would be quickly removed from the population, resulting in a dearth of detectable eQTLs (Ronald and Akey 2007). To test this, we compared the fractions of essential genes and nonessential genes among our eQTLs, while controlling for expression level (see Supplementary Text). We observed a slightly lower number of eQTLs in essential genes (permutation p = 0.036, Figure 3B), which becomes more pronounced for stronger eQTLs (permutation p = 4×10^−4^, Figure 3B). Similarly, we found that genes with more constrained protein sequences (as measured by the dN/dS ratio (Connallon and Knowles 2007)) also tend to have fewer eQTLs (permutation p = 2×10^−4^, Figure 3C), suggesting a congruence in purifying selection on expression levels and protein sequences.

### Molecular differences between clinical and non-clinical isolates

Although our analysis above confirmed that eQTLs are generally under negative selection, previous work has found that yeast cis-regulation can also be the source of adaptations (Chang et al. 2013, Naranjo et al. 2015). To explore this possibility, we next investigated whether any eQTLs have been subject to different evolutionary pressures between clinical and non-clinical isolates. *S. cerevisiae* is generally regarded as safe, but there have been case reports of invasive infection by *S. cerevisiae* across a wide range of patient types and locations (Enache-Angoulvant and Hennequin 2005; Munoz et al. 2005). These infections range from fungemia to endocarditis and are associated with conditions such as the placement of an intravenous catheter or an immune-compromised state. As a result, *S. cerevisiae* has been classified as an emerging opportunistic pathogen. This classification of a genetically tractable organism presents a unique opportunity for analyzing the evolutionary adaptations associated with pathogenicity. Gene expression adaptation promoting a pathogenic phenotype has been previous studied in *S. cerevisiae,* though the study focused on gene expression differences between only two isolates: BY4716, a laboratory strain, and YJM789, a derivative of a clinical strain (Fraser et al. 2012). With a larger number of strains in this study, we aimed to identify gene expression adaptation across a broader scope of clinical and non-clinical strains.

We first conducted a GWAS to identify genetic variants associated with the clinical niche. Muller *et al.* previously performed a GWAS using these same strains, with variants detected by a tiling microarray (Muller et al. 2011). Because the microarray could not precisely locate variants or distinguish multiple nearby variants, we reasoned that a GWAS using full genome sequences may have greater power due to lower genotyping error. Prior to performing the GWAS, we investigated the statistical power to detect associations given the population structure. Although population structure correction is prudent in GWAS of any species, this is particularly important in *S. cerevisiae* because of widespread admixture between strains (Connelly and Akey 2012). Using simulations of the population structure specific to our strains, we found that we have power to detect associations across a range of effect sizes (Supplemental Figure 15, Materials and Methods: “Genome-wide association testing”).

Our GWAS identified two SNPs reaching genome-wide significance (Figure 4A, p < 1.1×10^−6^, corresponding to FDR<0.05, Materials and Methods). One SNP (chromosome XII, position 830378) exhibited a particularly strong association (p = 2.9×10^−12^). The associated SNP is located 14 bp upstream of the ORF for *NIT3,* suggesting a cis-regulatory effect, and the lack of nearby associated variants (Figure 4B) suggests that this SNP is likely to be the causal variant. To test whether this variant could possibly be caused by sequencing error, we examined its read coverage and local LD structure; both analyses confirmed this is a high-confidence variant (Supplementary Text, Supplemental Figures 16 and 17). This variant was not previously detected, perhaps due to the lower resolution or accuracy of tiling array-based genotyping (Muller et al. 2011). Investigation of the population variation of this SNP suggests that the variant likely originated from a single mutation, as opposed to multiple independent mutations (Supplementary Text).

**Figure 4:**
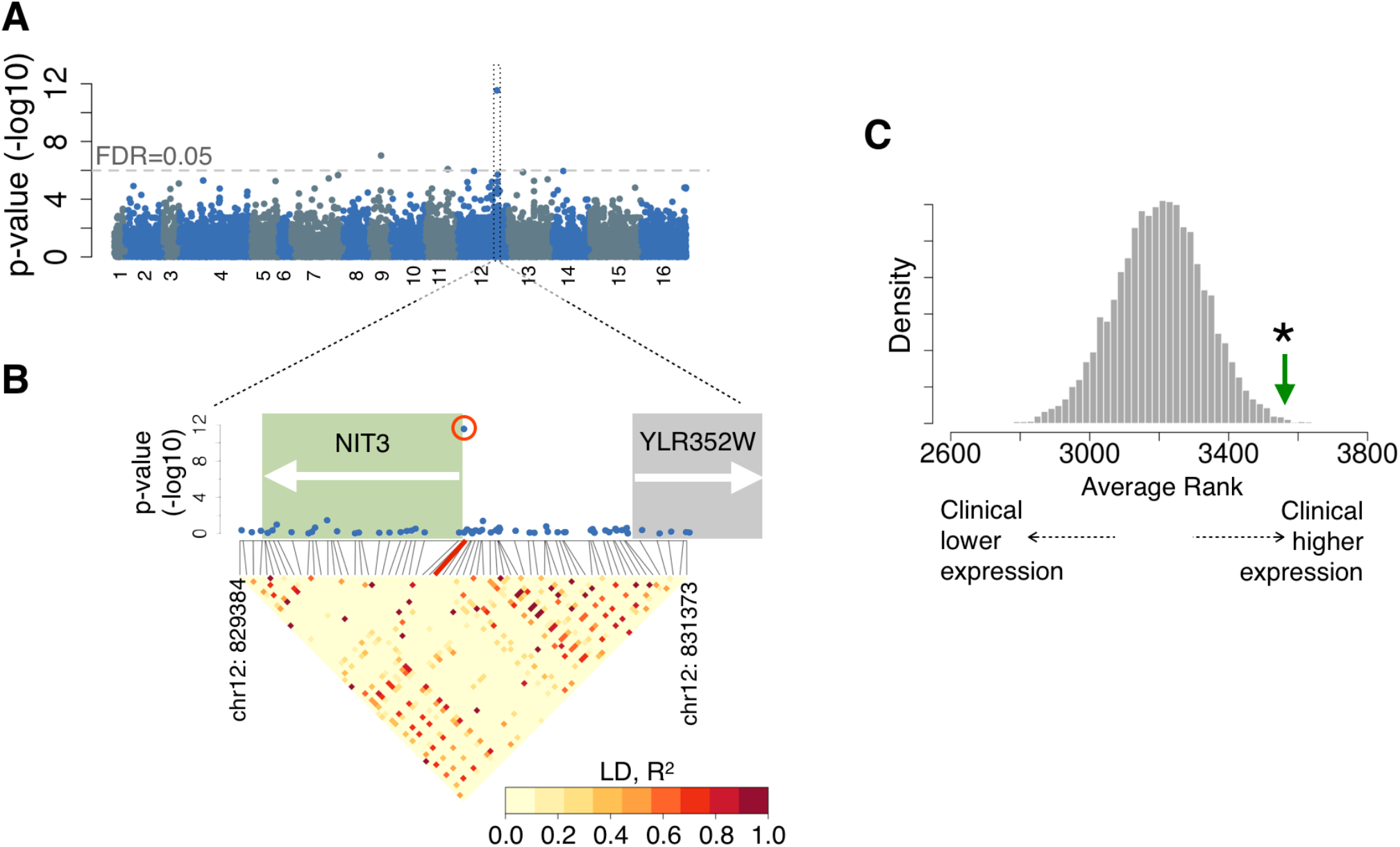
GWAS and differential expression of biofilm genes between clinical and non-clinical strains. (A) Manhattan plot of clinical and non-clinical variants. (B) Higher detail of the region neighboring the strongest GWAS variant. The locations of genes are shown behind the Manhattan plot and the linkage disequilibrium is shown below. (C) Expression of biofilm suppressor gene set. X-axis is the average rank of differential expression across the gene set. The gene with the most significant lower clinical expression has a rank of zero. Grey histogram presents the distribution from null permutations of the average rank from gene sets of the same size. The green arrow indicates the observed average rank.

*NIT3* is a member of the nitrilase superfamily, which encodes deaminating enzymes that play a role in biosynthesis of products such as biotin and auxin (Pace and Brenner 2001). The null mutant exhibits increased formation of biofilms (Vandenbosch et al. 2013), which are aggregate communities of adherent cells. Thus we classified the gene as a biofilm suppressor. Biofilm formation has been associated with virulence in *Candida albicans* and other fungi (Fanning and Mitchell 2012; Nobile and Johnson 2015). In *S. cerevisiae,* biofilm formation is assessed by the ability to adhere to plastic, and biofilm formation is associated with decreased growth rate, increased drug resistance, and a “mat”-like appearance (Reynolds and Fink 2001; Bojsen et al. 2014).

We next investigated whether gene expression differs between clinical and non-clinical strains using DESeq2 (Love et al. 2014) (Supplemental Figure 19). At an FDR of 0.05, we identified 325 differentially expressed genes (Supplemental Table 3). Because *NIT3* is a biofilm suppressor, we asked whether biofilm suppressors as a group exhibited differential expression between the clinical and non-clinical strains. 197 genes, including *NIT3,* were classified as biofilm suppressors because the deletion of the gene resulted in increased biofilm formation (Vandenbosch et al. 2013; Scherz et al. 2014). Comparing normalized expression levels between the clinical and non-clinical strains, we observed significantly increased expression of biofilm suppressors in the clinical strains (Mann-Whitney p=1.7×10^−5^), although *NIT3* was not differentially expressed. Because this observation could be driven by a small subset of genes, we also performed a more conservative test. For this test, we first ranked all genes based on their significance and directionality of differential expression between clinical vs. non-clinical strains. We then tested whether the average rank of biofilm suppressors was significantly different than the average rank of random sets of the same number of genes. Again, biofilm suppressors were significantly overexpressed in the clinical strains (permutation p = 1.9×10^−3^, Figure 4C). The increased expression of the biofilm suppressors in the clinical strains suggests that the clinical strains have an overall gene expression profile consistent with disruption of biofilm formation.

### Evidence for lineage-specific selection on biofilm suppressors

We then proceeded to test whether this expression difference in biofilm suppressors was driven by natural selection acting on eQTLs. This question is important because although we found an overall gene expression difference in biofilm suppressors, this is not by itself an indication of gene expression adaptation. By chance, neutral drift of even a single variant (such as a mutation in a transcription factor) could single-handedly account for the differential expression of many genes. One approach to surmount this challenge is to identify an excess of independent regulatory variants that act in the same direction (e.g. up-regulation in clinical isolates), since this would not be expected under neutral evolution (Fraser 2011). This sign test approach has been used to identify several examples of polygenic adaptation in yeast and other species (see Introduction).

For each eQTL, we assessed whether the clinical strains exhibited a general up- or down-regulation of the gene by calculating the difference in the up-regulating allele frequency among the clinical strains and the non-clinical strains (Figure 5A,B). An overall positive value indicates that the clinical strains had a higher up-regulating allele frequency, suggesting that the local genetic control of that gene is shifted toward up-regulation in the clinical strains. The eQTLs with the largest positive and negative allele frequency differences are shown as examples (Figure 5C). Interestingly, both genes are involved in endocytosis, which is a function that has been previously identified to be under lineage-specific selection in clinical strains (Fraser et al. 2012).

**Figure 5:**
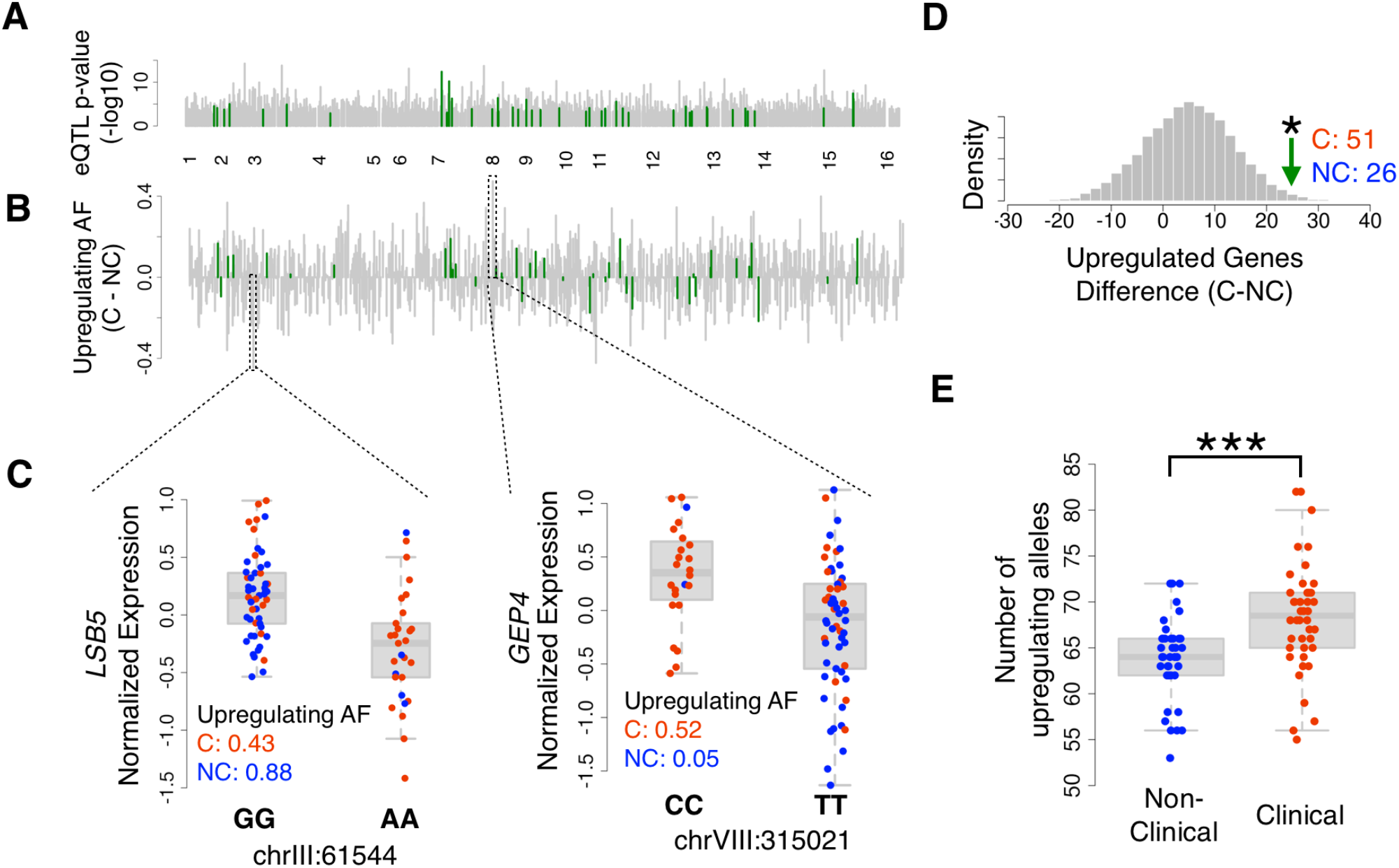
Testing for adaptation in biofilm suppressors. (A) Locations of cis-eQTLs across the genome (FDR<0.05). The biofilm suppressors that have significant cis-eQTLs are highlighted in green. (B) Difference in upregulating allele frequency between the clinical and non-clinical strains for each of the cis-eQTLs. (C) Examples of cis-eQTLs with the largest positive and negative difference in upregulating allele frequencies. Each point represents a strain, with the color indicating whether the strain is clinical (red) or non-clinical (blue). (D) Permutation test to assess the significance of the number of positive upregulating allele frequencies in the clinical strains. The x-axis indicates the difference in the number of genes that have higher upregulating allele frequencies in the clinical strains and the number of genes that have higher up-regulating allele frequencies in the non-clinical strains. The green arrow marks the difference found in biofilm suppressors, and the histogram indicates the difference found across each of 1000 permutations. (E) The number of up-regulating alleles for each strain segregated by clinical and non-clinical origin.

Using these directionalities, we next investigated the overall directionality of the local eQTLs in biofilm suppressors. To obtain a larger number of biofilm genes with local eQTLs, we used a less stringent cutoff to define eQTLs (empirical p < 0.2, which is the least stringent cutoff at which there is significant concordance in directionality with the Skelly *et al.* dataset; Supplemental Figure 10). Among the biofilm suppressor genes, we identified 51 local eQTLs with higher up-regulating frequency in the clinical strains, and 26 local eQTLs with higher up-regulating frequency in the non-clinical strains. To assess the significance of this difference, we repeated the analysis with 10,000 random sets of 77 local eQTLs. This permutation procedure accounts for population structure and demographic confounders because the difference within the biofilm suppressors is compared with the rest of the genome (method adapted from Fraser 2013). Using this test, we found significantly more clinical-upregulated biofilm suppressors than expected (permutation p = 7.3×10^−3^, Figure 5D). The occurrence of this many independent regulatory variants shifting in the same direction (up-regulation in clinical isolates) is unlikely to occur under neutral evolution. Analogous to the sign test in allele-specific expression analyses, this indicates that the biofilm suppressors are likely under lineage-specific selection. Since human infection is almost certainly not the ancestral niche of *S. cerevisiae*, we can additionally infer that the change is most likely due to up-regulation in response to novel selection pressures on the clinical strains, rather than down-regulation in non-clinical strains.

We next assessed whether the allele frequency difference was occurring in a small sample of clinical strains, or if this was a general trend across all clinical strains. To investigate this distribution, we counted the number of up-regulating biofilm eQTL alleles for each strain (Figure 5E). We observed a general shift in distribution suggesting that the overall up-regulation occurred as a trend across the majority of clinical strains (Mann-Whitney p = 1.7×10^−4^). Concordant with the differential expression analysis, these results provide evidence that the clinical strains have undergone adaptation to increase expression of the biofilm suppressors.

## DISCUSSION

In this work, we identified local eQTLs regulating ~20% of the genes in *S. cerevisiae.* Global eQTL mapping has been performed in *S. cerevisiae* before, using crosses between two strains (Brem et al. 2005, Androset et al. 2011, Gagneur et al. 2013) or a set of 22 isolates (Skelly et al. 2013). Although these previous eQTL studies have revealed rich information on the genetics of gene expression, mapping with 85 diverse strains has the advantages of greater genetic diversity and less linkage disequilibrium between nearby variants. These qualities facilitated fine-mapping of causal eQTL variants and also allowed us to explore the species-wide evolution of eQTLs.

Assessing all of the eQTLs, we found that their 1-LOD support intervals are ~50-fold smaller than previous yeast eQTLs, and that they are enriched near the transcription start and end sites of genes. Previous studies in several species have found eQTL enrichment near transcription start and end sites (Veyrieras et al. 2008; Stranger et al. 2012; Veyrieras et al. 2012; Huang et al. 2015; Tung et al. 2015), as well as higher genetic variation in 3’ UTRs of genes with allele-specific expression (Ronald et al. 2005); however to our knowledge our eQTLs represent the first example of eQTLs implicating a specific class of binding sites within 3’ UTRs. We found that the 3’ UTR eQTLs are enriched for RNA-binding protein motifs, suggesting that a common mechanism for these eQTLs is post-transcriptional. We did not find any evidence for eQTLs within ORFs to act in *trans,* since there was no enrichment for nonsynonymous variants that would be much more likely to mediate *trans*-acting effects. It is important to note that in this study, we identified the location of the eQTL relative to the most common isoform for each gene. Previous analysis has found a remarkable diversity in isoforms in yeast (Pelechano et al. 2013), and thus future analyses on the association between isoform diversity and eQTL location would be informative.

We then used these high-resolution eQTLs to study the evolution of gene expression across our 85 strains. Previous studies in several species have shown that gene expression is predominantly under stabilizing selection, e.g. in mutation accumulation experiments where organisms are grown for many generations with minimal selection (Denver et al. 2005; Rifkin et al. 2005). Indeed, we confirmed that eQTLs are generally under purifying selection (Ronald and Akey 2007, Battle et al. 2014; Josephs et al. 2015), and that eQTLs are depleted among essential genes and evolutionarily constrained genes (Ronald and Akey 2007).

Despite the predominance of stabilizing selection, there is growing evidence that positive selection is also an widespread force acting on regulatory variation in a wide range of species (Fraser et al. 2011; Fraser 2013; Enard et al. 2014). To search for gene expression adaptation in these strains, we tested for lineage-specific selection on eQTLs between the clinical and non-clinical strains. Our GWAS and differential expression analysis independently implicated biofilm suppressors as a differentiating gene set. Consistent with this, we found that allele frequency shifts of eQTLs have led to increased expression of biofilm suppressor genes in clinical strains. Although the allele frequency shifts were generally small, by testing across a large number of genes, we were able to identify a significant effect that is not consistent with neutral evolution (Fraser 2011).

Biofilm regulation has been previously studied in other medically-relevant fungi (Fanning and Mitchell 2012), and has a particularly rich literature in *Candida albicans* (Nobile and Johnson 2015). Although *Candida* and *Saccharomyces* are evolutionarily distant, elements of the biofilm formation pathways are conserved between them (Nobile et al. 2012). Notably, we found evidence for increased biofilm suppression in the clinical strains, which is contrary to the standard impression of biofilm activation in pathogenic microbes. We hypothesize several possible explanations: (1) biofilm dispersal is a key component of the biofilm virulence effect (Uppuluri et al. 2010; Nobile and Johnson 2015; Uppuluri and Lopez-Ribot 2016), and the increased expression of biofilm suppressors may promote this dispersal; or (2) the classification of biofilm genes as general suppressors or activators may be too simplistic—e.g. ignoring condition-dependent effects—and thus, our results could reflect adaptation in a specific but uncharacterized component of the biofilm process. Furthermore, these genes are likely to have a variety of functions unrelated to biofilms, and thus the effects that we observed may even represent selection acting on a different phenotype. Future experiments will be essential in determining whether these eQTLs lead to increased pathogenicity, or affect some other aspect of adaptation to human hosts.

In addition to further investigating the role of biofilm formation, future studies may reveal much more about these strains. For example, recent studies have found a large number of ploidy and copy number variations between strains of *S. cerevisiae* (Dunn et al. 2012; Bergstrom et al. 2014). In this study, although the absence and presence of genes were assessed, we did not estimate copy number (see Supplementary Text). Instead, by measuring gene expression, we were able to directly measure the amount of mRNA and thus measure the downstream effects of whatever copy number variation exists. The difference, however, between gene expression regulation by eQTL versus copy number variation remains to be investigated. Another area of future study is the measurement of gene expression in multiple conditions, since eQTLs can be environment-specific (Smith and Kruglyak 2008). In addition, work in *C. albicans* has found that gene regulatory relationships differ between infection environments and laboratory media (Xu et al. 2015). Such environment-specific expression might explain the lack of differential expression seen in *NIT3* (though this variant might also act at another level of regulation, such as translation initiation, another major source of divergence in yeast gene regulation (Artieri and Fraser 2014; McManus et al. 2014)).

The strength of *S. cerevisiae* as a model organism is based upon its easy manipulation, well-scrutinized genes, and a rich landscape of strains across diverse geographical and ecological niches. In this study, we contribute to this compendium of knowledge with genome-wide gene expression across many strains and a high-resolution map of genetic regulation of gene expression. We have shown that the gene expression across these strains is not only constrained, but also under pressure to adapt across the varied strain histories and phenotypes. We hope for this resource to serve as the launching point for future studies on the mechanisms and consequences of gene expression variation.

## METHODS

### Strain selection and DNA sequencing

85 isolates of *S. cerevisiae* were obtained from the Pfaff Yeast Culture Collection (Supplementary Table 1). Locations of isolation were obtained from Muller *et al.* 2011. Isolates were grown overnight in YPD liquid culture at 30°C, and DNA extraction was performed using the MasterPure Yeast DNA purification kit. Pooled, barcoded libraries were prepared using the Nextera Sample Preparation Kit. Sequencing was performed using Illumina Hi-Seq 2000 with paired-end 101 base-pair reads. Reads were then dynamically trimmed and length-sorted using SolexaQA default parameters (Cox et al. 2010). Mapping was performed using STAMPY with default parameters and substitution rate 0.005 (Lunter and Goodson 2011). Reads were then sorted, and duplicates were removed using Samtools (Li et al. 2009). Variant calling was performed using freeBayes (default parameters, T=0.005) using all BAM files together (Garrison and Marth 2012). To compare the percent concordance of the allele calls in the strains that were sequenced in both this study and Strope *et al.* 2015, we also performed mapping and variant calling on the Strope dataset using the same parameters. For downstream analysis using GEMMA, which requires no missing variants, we removed SNPs with greater than 10% missing data and imputed the rest of the missing variants using MACH (--mle --rounds 100 --states 200) (Li et al. 2010).

### RNA sequencing

Strains were grown to log-phase growth in YPD at 30°C, and total RNA was extracted using the MasterPure Kit. Sequencing libraries were constructed using the Illumina TruSeq kit. 36 bp-single end reads were sequenced using the Illumina HiSeq 2000 across eight lanes with a randomized lane assignment to prevent batch effects. The reads were dynamically trimmed using DynamicTrim and LengthSort of the SolexaQA package with default parameters (Cox et al. 2010). Reads were then mapped using STAR version 2.4.1 to the S288c reference genome with all SNPs from the DNA-sequencing analysis masked (Dobin et al. 2013). The genome index was generated with flag “--genomeSAindexNbases 11” and mapping was performed with default parameters. STAR mitigates mapping bias by allowing for mismatches. The lack of genome-wide mapping bias was confirmed by a lack of significant correlation between percent reads mapping and the genomic distance from the S288c reference genome (Supplemental Figure 4). Principal components of the expression data were calculated using the standardized RPKM expression, excluding genes with a sum of RPKM across all strains < 1. In addition, we assessed whether genes were missing using the DNA sequencing data, and these genes were marked as missing for subsequent eQTL mapping (Supplementary Text).

### eQTL mapping

RPKM values were quantile-normalized across strains and the resulting values were fit to a standard normal for each gene. We then used PEER to discover hidden covariates (k=15), and these covariates were regressed out (Stegle et al. 2010). The removal of hidden covariates improves power in cis-eQTL mapping and also can remove unwanted batch effects (Stegle et al. 2010, Leek and Story 2007, Johnson et al. 2007). We performed associations with variants with minor allele frequency greater than 0.1 using GEMMA (Zhou and Stephens 2012). Because GEMMA does not allow missing variants, we imputed the variants using MACH 1.0 (Li et al. 2010), removing any variants with more than 10% missing. The relatedness matrix for association testing was calculated using all variants after LD pruning (r^2^ > 0.8, sliding window) with the centered relatedness approach, as described in the GEMMA manual. To assess significance, we performed 1,000 to 10,000 permutations, comparing for each gene, the best-associating p-value from each permutations with the best-associating p-value from the non-permuted associations. The permutations provided empirical p-values, from which we assessed false discovery rates using the Benjamini-Hochberg method (Benjamini and Hochberg 1995). We compared results from these permutations performed on each transcript individually with that of permutations performed preserving the relationship across transcripts - which revealed similar empirical p-values (Supplemental Figure 11). To localize the eQTLs relative to genic components, we used annotations of open reading frames from SGD (Cherry et al. 2012). In cases where multiple eQTLs for the same gene were tied for most significant, the midpoint between the minimum and maximum position was used to represent the eQTL location. Transcript boundaries of the isoform with the highest number of supporting reads were used (Pelechano et al. 2013). Synonymous vs non-synonymous variants were identified with SNPeff (Cingolani et al. 2012).

### Genome-wide association testing

Genome-wide association testing was performed using variants with no missing genotypes, MAF > 0.05, and LD pruned by PLINK (r^2^ > 0.8, sliding window) (Purcell et al. 2007; Chang et al. 2015). To identify the most effective method for mapping between genotypes and traits, we performed a power analysis using three methods (Supplementary Text). We found that GEMMA performed best at controlling for false-positives due to population structure while maintaining power. We thus performed all subsequent association testing with GEMMA. The relatedness matrix was calculated using all tested variants with the centered relatedness approach, as described in the GEMMA manual. Calculating significance after GWAS used a simpler approach than eQTL mapping because all variants were tested together rather than using a gene-by-gene approach. The cutoff for false discovery rate was identified by 10^5^ permutations of a randomized phenotype, identifying 1.02x10^−6^ as the cutoff for FDR < 0.05.

## Data

All RNA-Seq and DNA-Seq data are deposited in Bioproject PRJNA342356 in the NCBI Sequence Read Archive

## Acknowledgements

We would like to thank members of the Fraser lab for helpful discussions, and K. Boundy-Mills for providing yeast strains. This work was supported by NIH grant 2R01GM097171-05A1.

## SUPPLEMENTAL FIGURES

Section A: DNA and RNA-Seq of 85 yeast strains

Supplemental Figure 1: Assessment of the quality of the DNA-Sequencing
Supplemental Figure 2: Assessment of the concordance of variant calls with Strope et al.
Supplemental Figure 3: Phylogenetic tree close-up with ecological niche and strain names
Supplemental Figure 4: Assessment of RNA-Seq quality
Supplemental Figure 5: Assessment of absence of Genes
Supplemental Figure 6: Assessment of absent gene location
Supplemental Figure 7: Principal components of gene expression
Supplemental Figure 8: Assessment of batch effects on gene expression
Supplemental Figure 9: Association between ecological niche and principal components of genotype.

Section B: High resolution local-eQTL mapping

Supplemental Figure 10: Concordance of eQTLs with Skelly et al.
Supplemental Figure 11: Comparison between relationship-preserved eQTL permutations
Supplemental Figure 12: Local eQTL resolution and effect sizes of eQTLs according to location.
Supplemental Figure 13: Homer enrichment result for the 3’ UTR eQTLs
Supplemental Figure 14: Homer enrichment result for the promoter regions

Section C: Global Selection on eQTLs

No supplemental figures.

Section D: Molecular differences between clinical and non-clinical strains

Supplemental Figure 15: Power analysis of GWAS and population-structure correction
Supplemental Figure 16: Assessment of genome-wide linkage disequilibrium.
Supplemental Figure 17: Assessment of the quality of the chrXI: 830378 SNP
Supplemental Figure 18: Regional neighbor-joining tree of region surrounding NIT3
Supplemental Figure 19: Differential expression between clinical and non-clinical strains

**Supplemental Figure 1:**
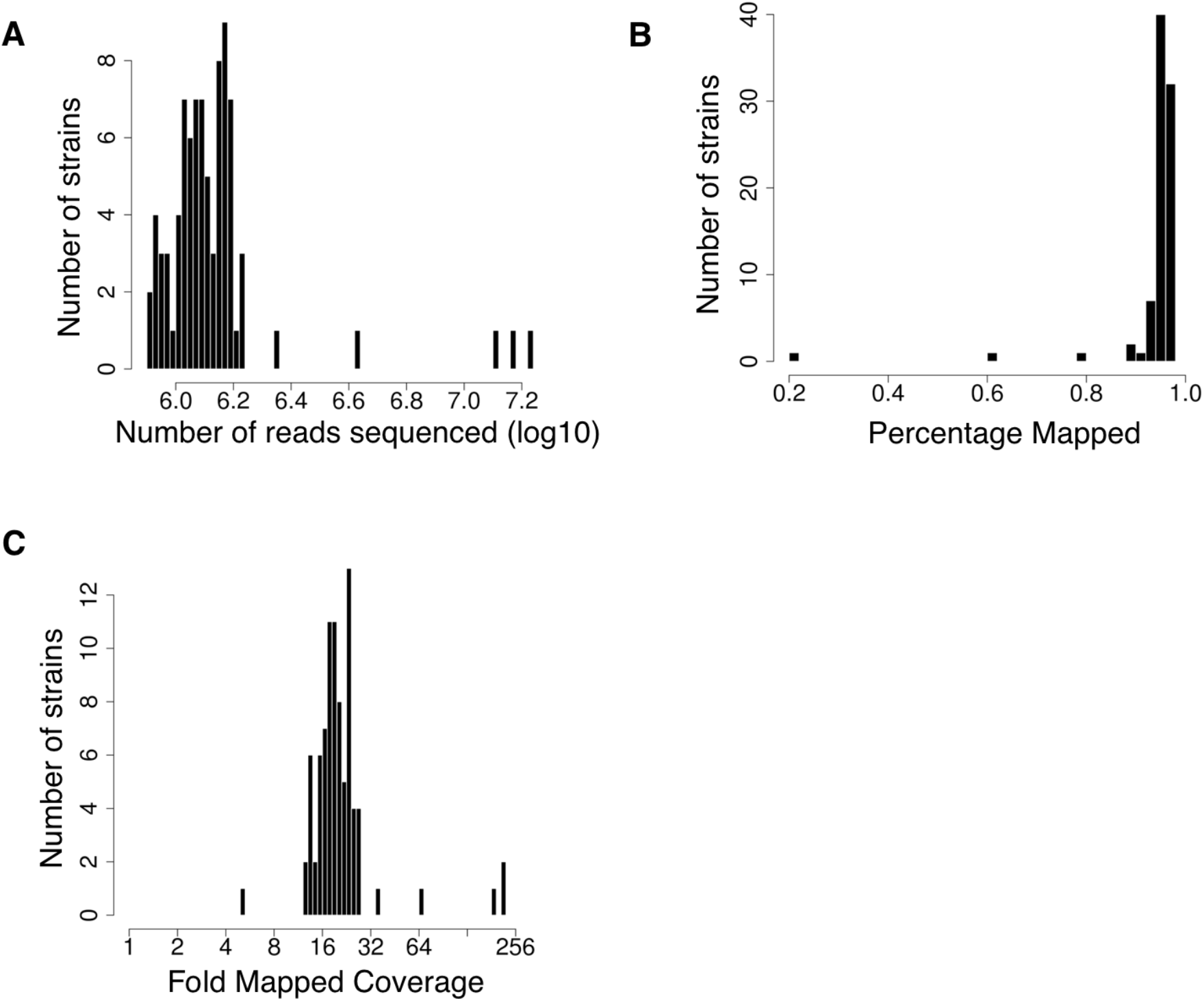
Assessment of the quality of the DNA-sequencing. (A) Histogram of the number of reads sequenced (log10) for each strain. (B) Histogram of the mapping percentage for each strain. (C) Histogram of the fold map coverage for each strain. Note that three of these strains (YJM981, YJM1307, YJM1400) have low mapping percentage. We justified keeping these strains for analysis based on two reasons. 1. Despite the low mapping percentage, the fold map coverage is still above 4 (see panel C). 2. These three strains were also sequenced by Strope et al. 2015 with strong concordance (Supplemental Figure 2).

**Supplemental Figure 2:**
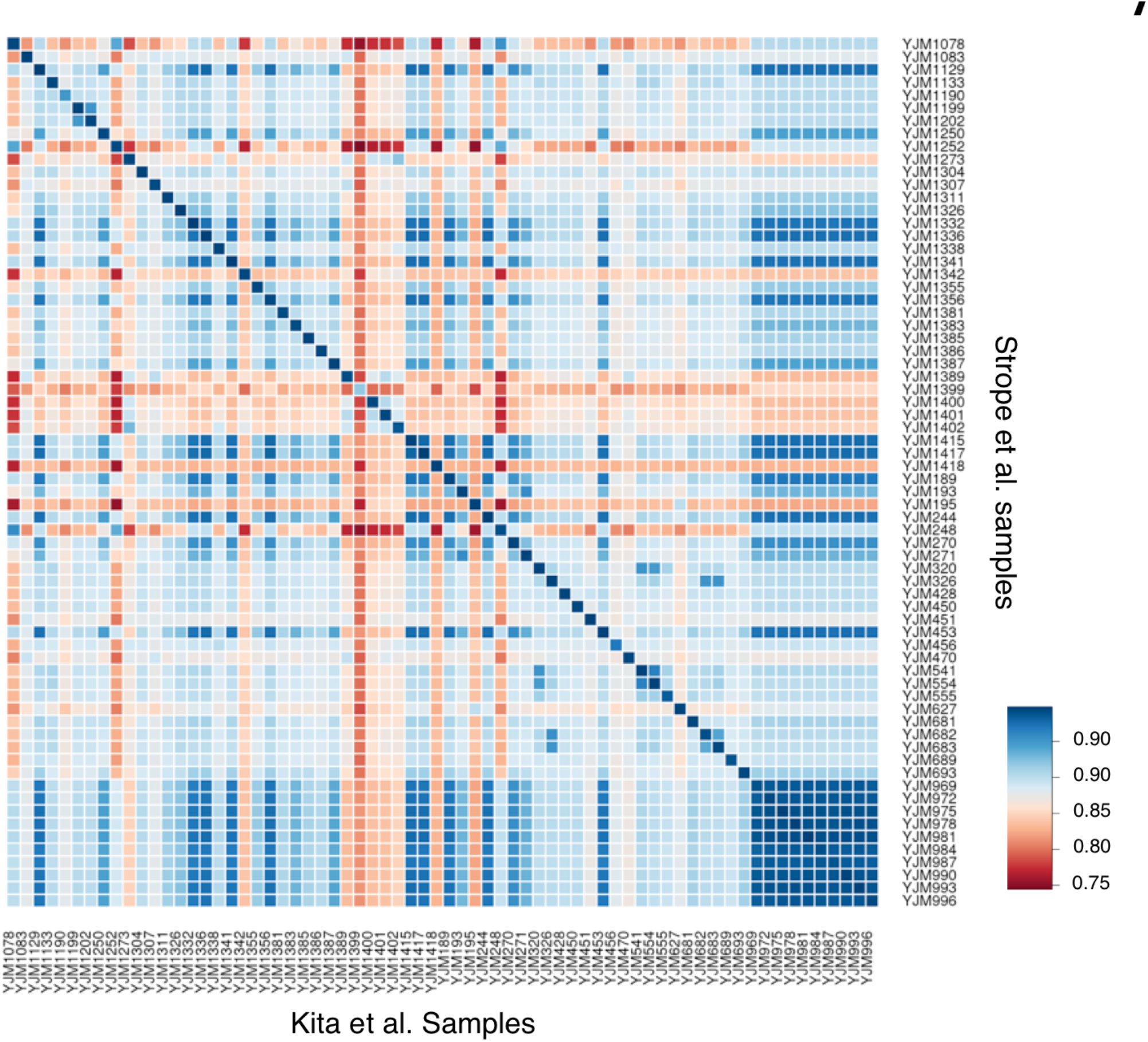
Assessment of the concordance of variant calls with Strope et al. Percent concordance of allele calls between every pair of strains (one from this study and one from Strope et al. 2015) were calculated across all variants common between the strains. Blue indicates high concordance and red indicates weak concordance.

**Supplemental Figure 3:**
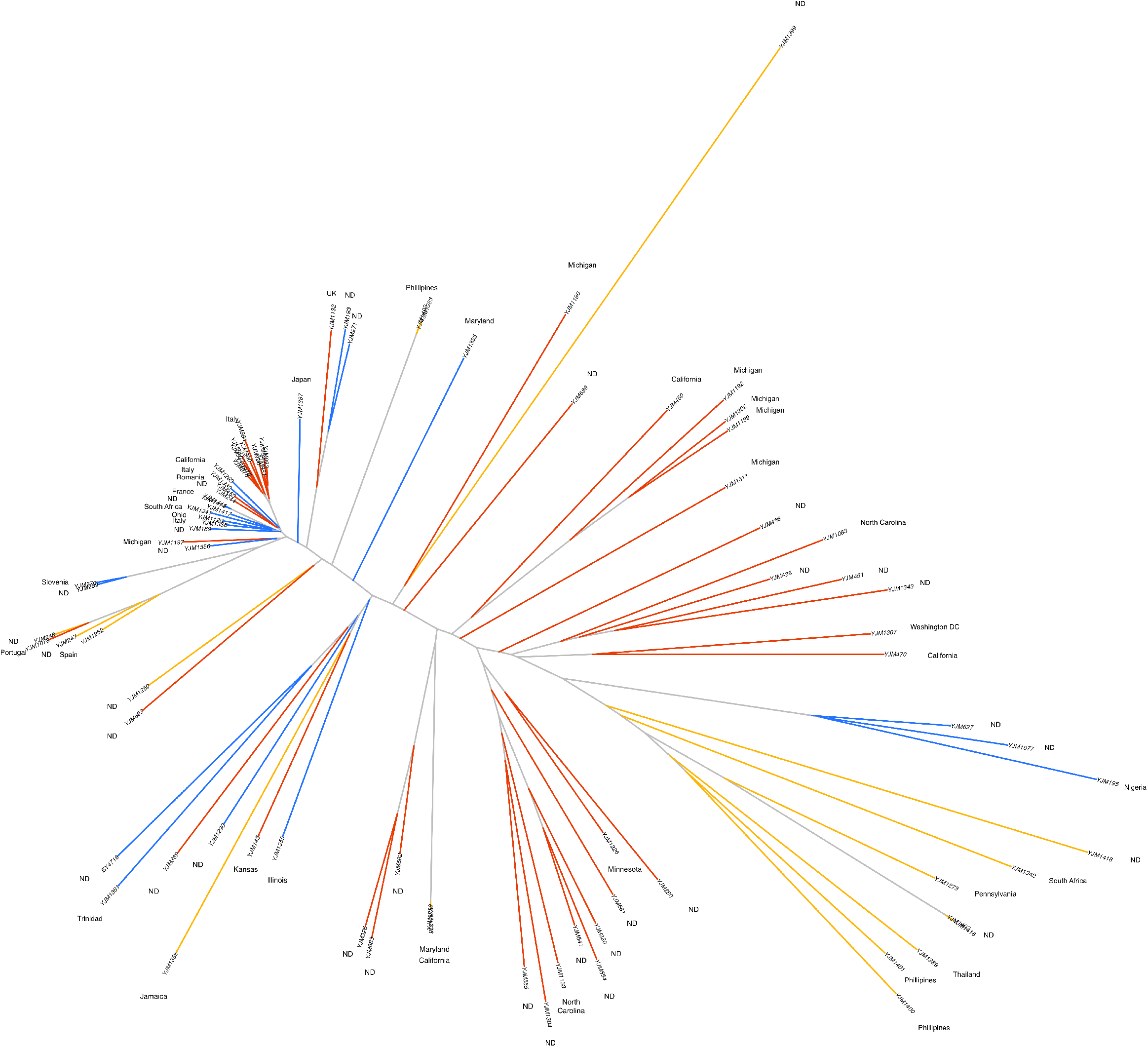
Phylogenetic tree close-up with ecological niche and strain names. An expanded figure of Figure 1B with the ecological niche and strain names. The colors for the ecological niche match Figure 1B.

**Supplemental Figure 4:**
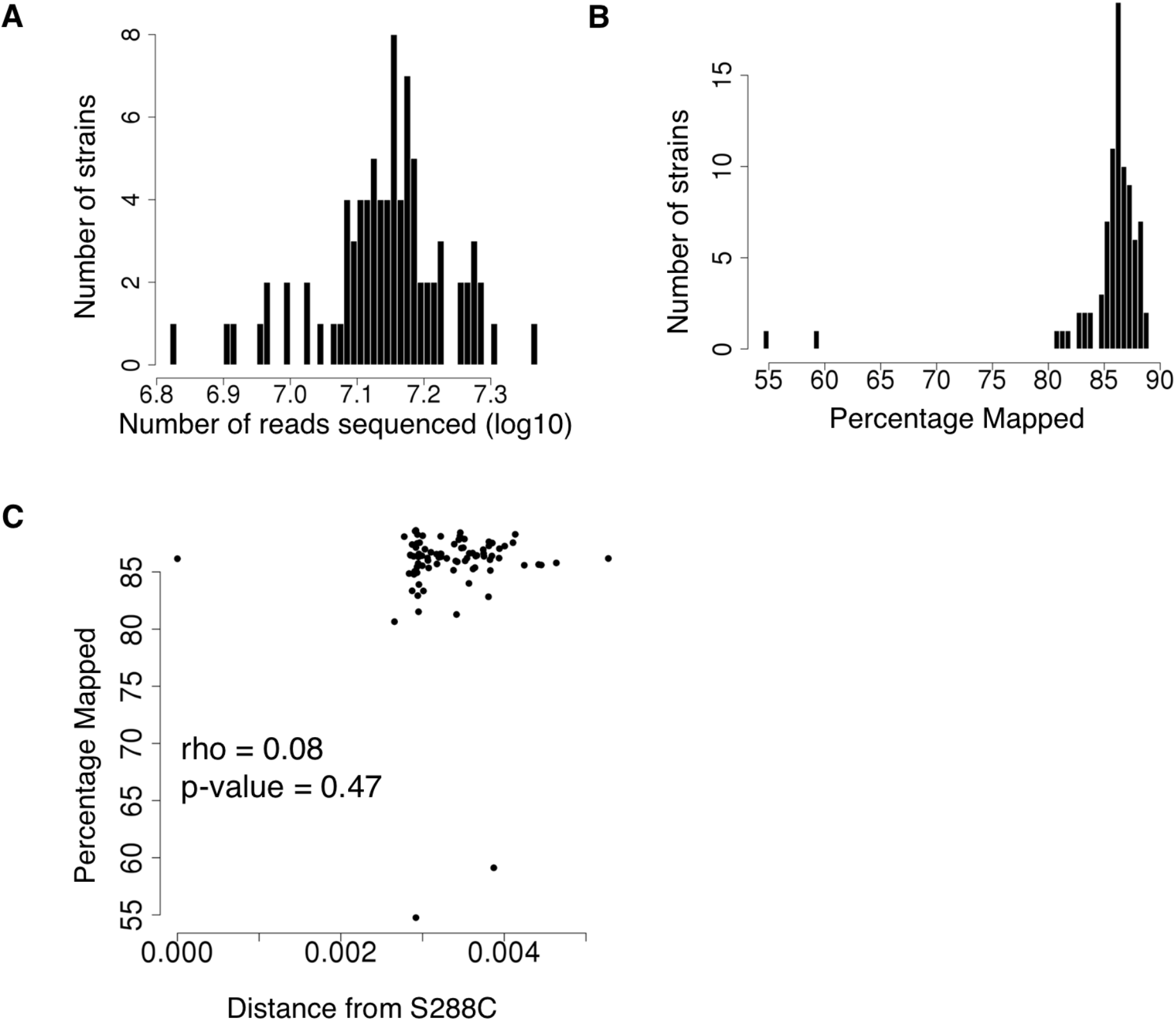
Assessment of RNA-Seq quality. (A) Histogram of the number of reads sequenced in each strain. (B) Histogram of the percent mapped reads in each strain. The low mapping rates in the two outlier strains is a result of high rates of nonunique mapping. (C) Evaluation of genome-wide mapping bias by associating the percentage of mapped reads with the genomic distance from S288c. Genomic distance was calculated as the percent variants that differed across the set of variants common to all strains. If mapping bias was present, we would expect that strains that are more distant from S288c would exhibit less mapping. We do not see this pattern as assessed by Spearman correlation.

**Supplemental Figure 5:**
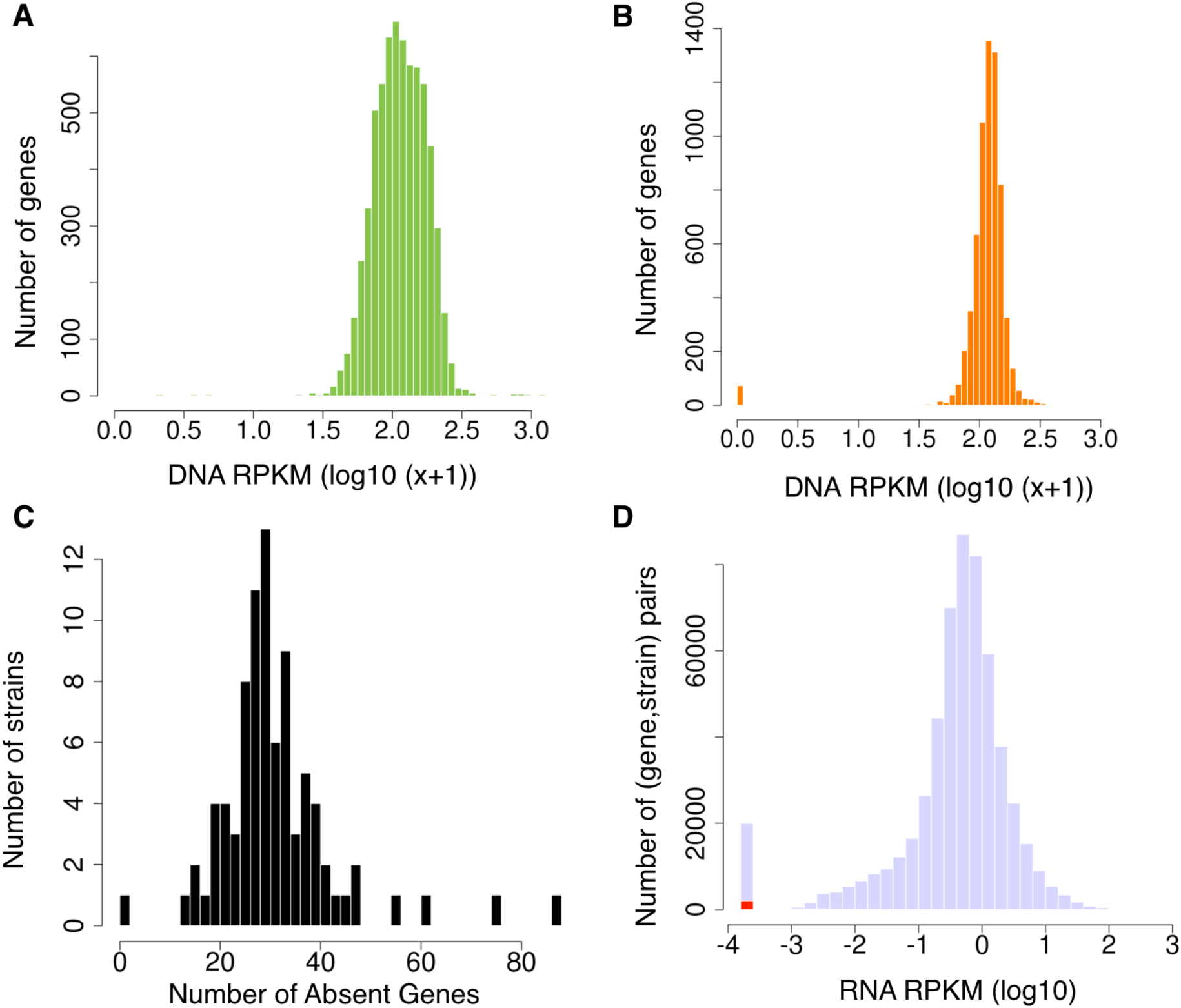
Assessment of absence of Genes. (A) Distribution of DNA-sequencing coverage over each gene for BY4716, the reference genome. (B) As an example, the distribution of DNA-sequencing coverage over each gene for YJM1418. (C) Distribution of number of absent genes for each strain sequenced in this study. (D) For each (gene, strain) pair the corresponding RNA-Seq expression was extracted. The histogram of the expression values is shown, where the (gene, strain) pairs that were marked as “absent” in red.

**Supplemental Figure 6:**
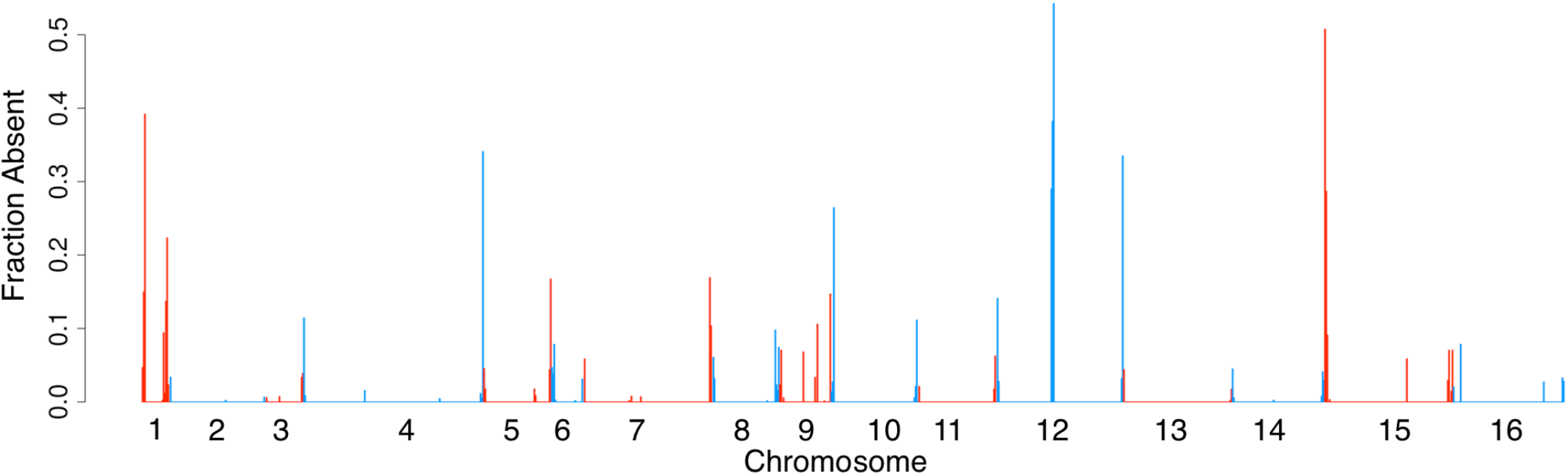
Assessment of absent gene location. Fraction absent genes per 10 KB window (gene position assigned to middle of its boundaries) aggregated across all strains. The fraction absent is calculated by (Sum of Number of absent genes in window across all strains) / (Total possible genes in window * Number of strains). For example, the peak near the end of chromosome 4 indicates that every gene within that 10 KB window was absent in all strains. The colors of consecutive chromosomes are alternated for visualization purposes.

**Supplemental Figure 7:**
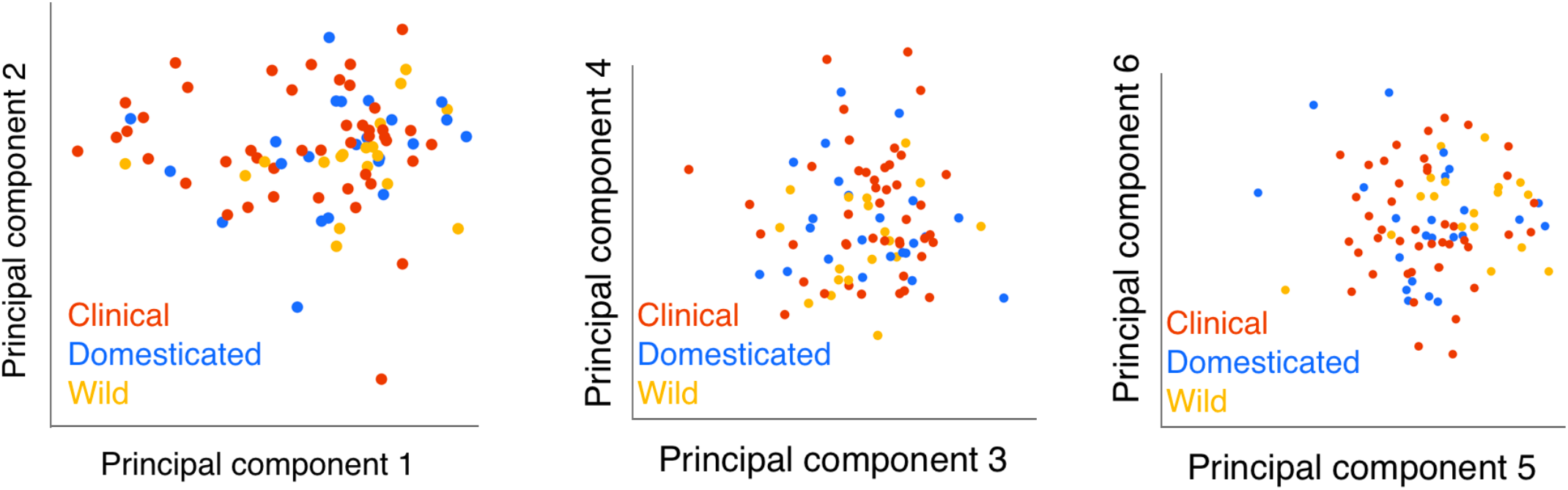
Principal components of gene expression. Principal components 1-6 of genome-wide gene expression. Note that the strains do not segregate by origin.

**Supplemental Figure 8:**
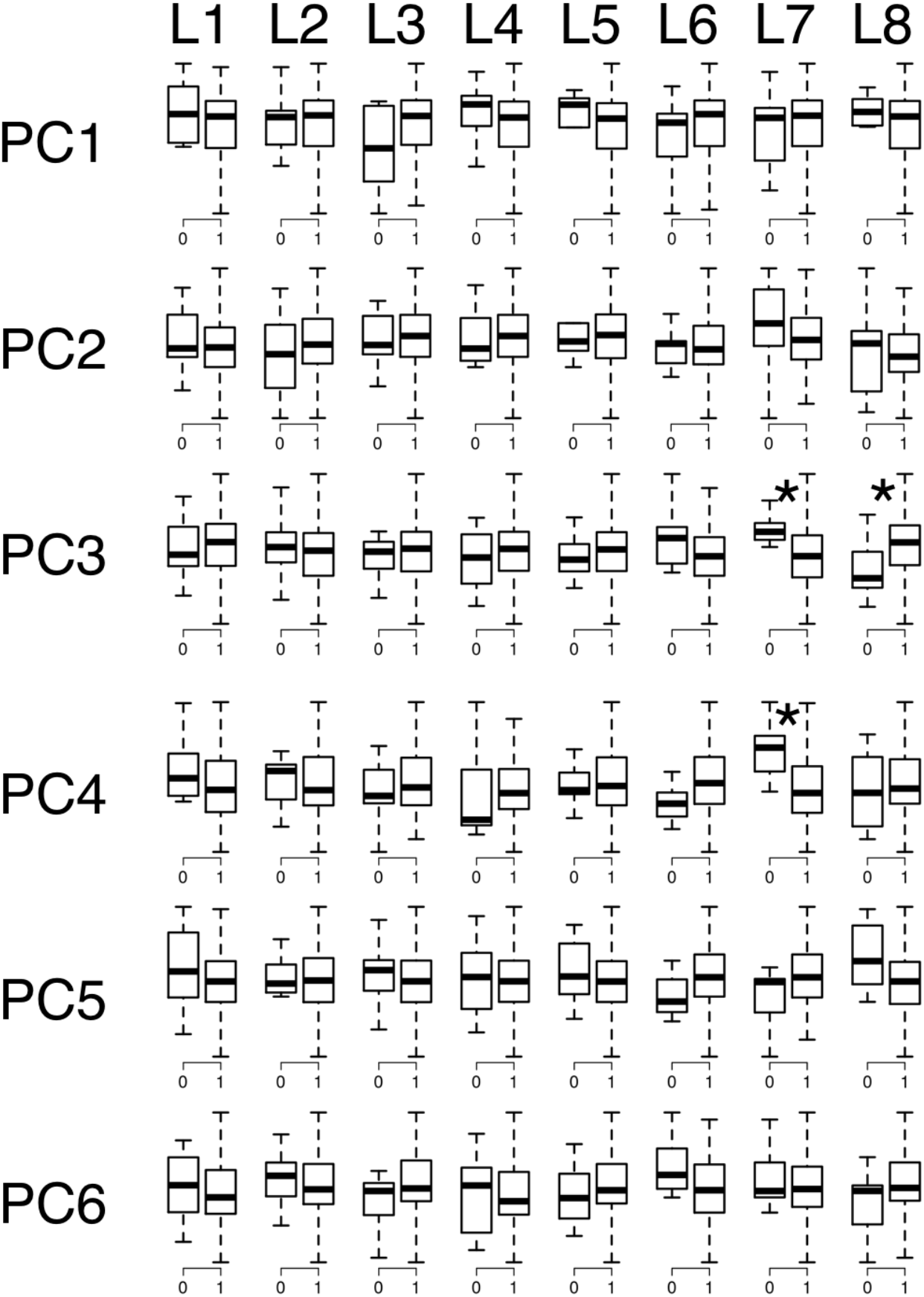
Assessment of batch effects on gene expression. The first six principal components were associated with each lane of sequencing. * indicates a nominal p-value < 0.05.

**Supplemental Figure 9:**
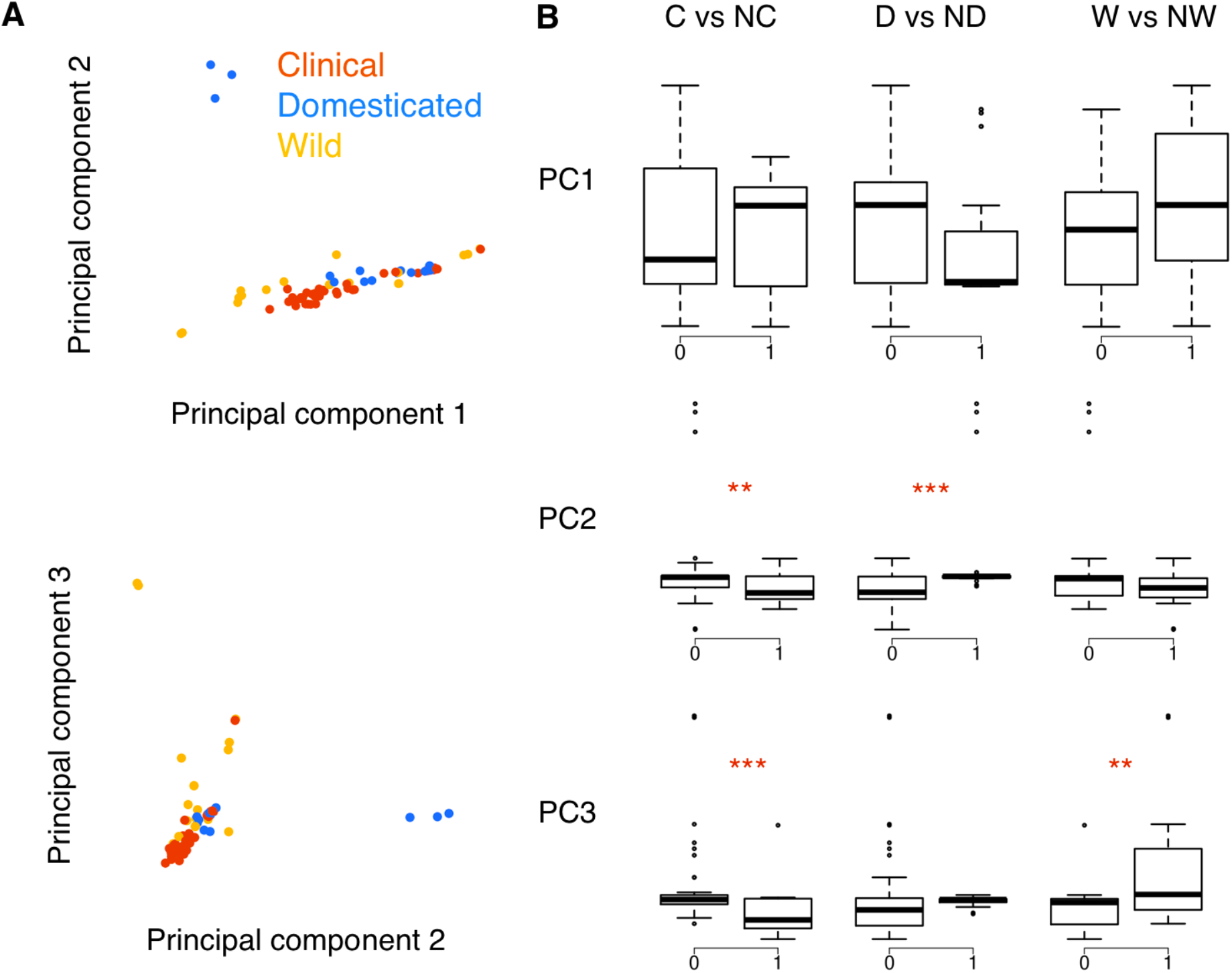
Association between ecological niche and principal components of genotype. A. The principal components were calculated from the conserved 218 kb sequence used to create the neighbor-joining trees. B. Boxplots comparing the principal components of each ecological niche category (C: clinical, NC: non-clinical, D: clinical, ND: non-clinical, W: wild, NW: non-wild. ** p-value < 0.005, *** p-value < 0.0005).

**Supplemental Figure 10:**
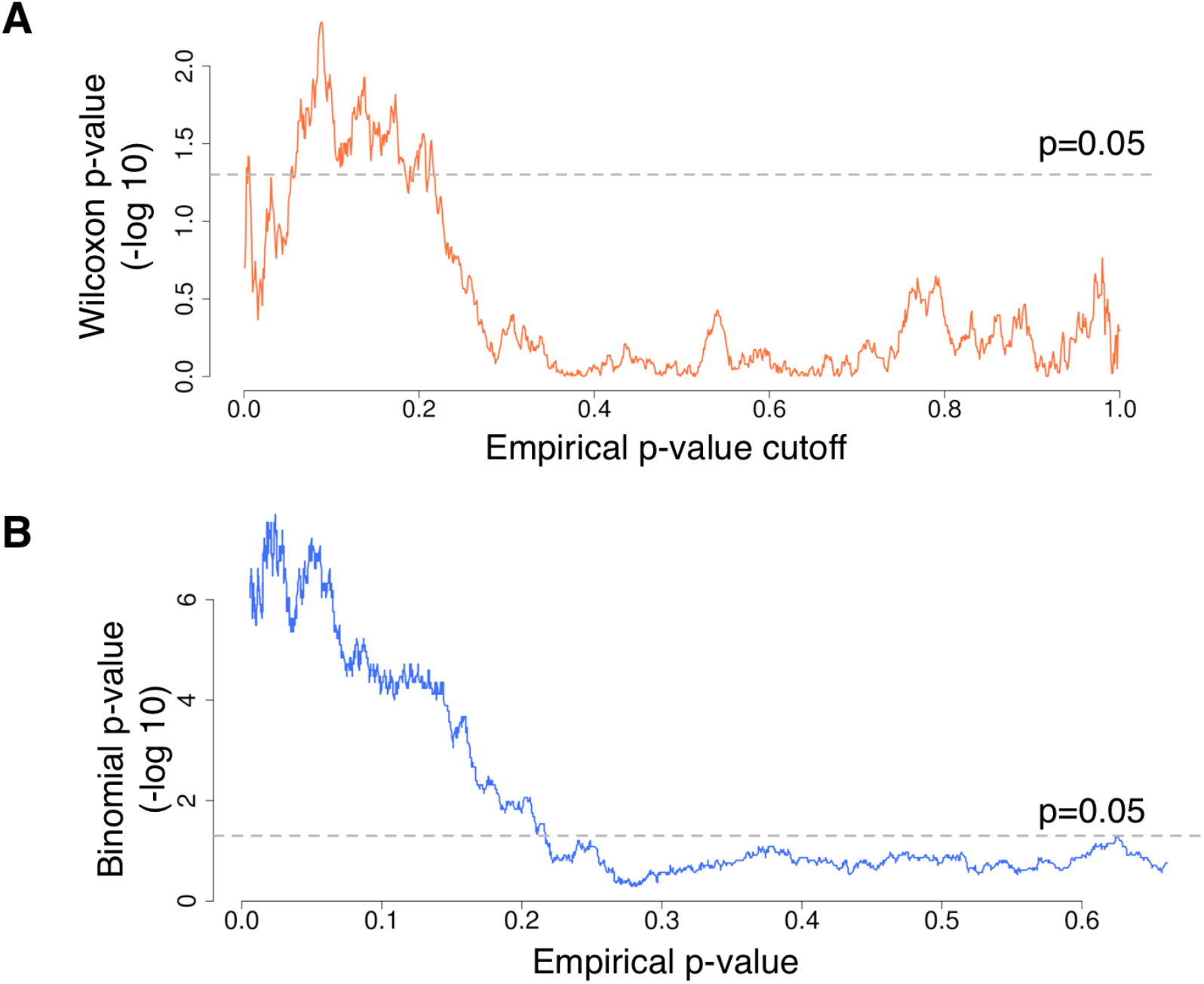
Concordance of eQTLs with Skelly et al. (A) Comparison of eQTL association p-values with Skelly *et al.* The x-axis indicates the empirical p-value cutoff. See Supplementary Text for details (B) Comparison of eQTL directionality with Skelly et al. The x-axis indicates the median empirical p-value of the sliding window. See Supplementary Text for details.

**Supplemental Figure 11:**
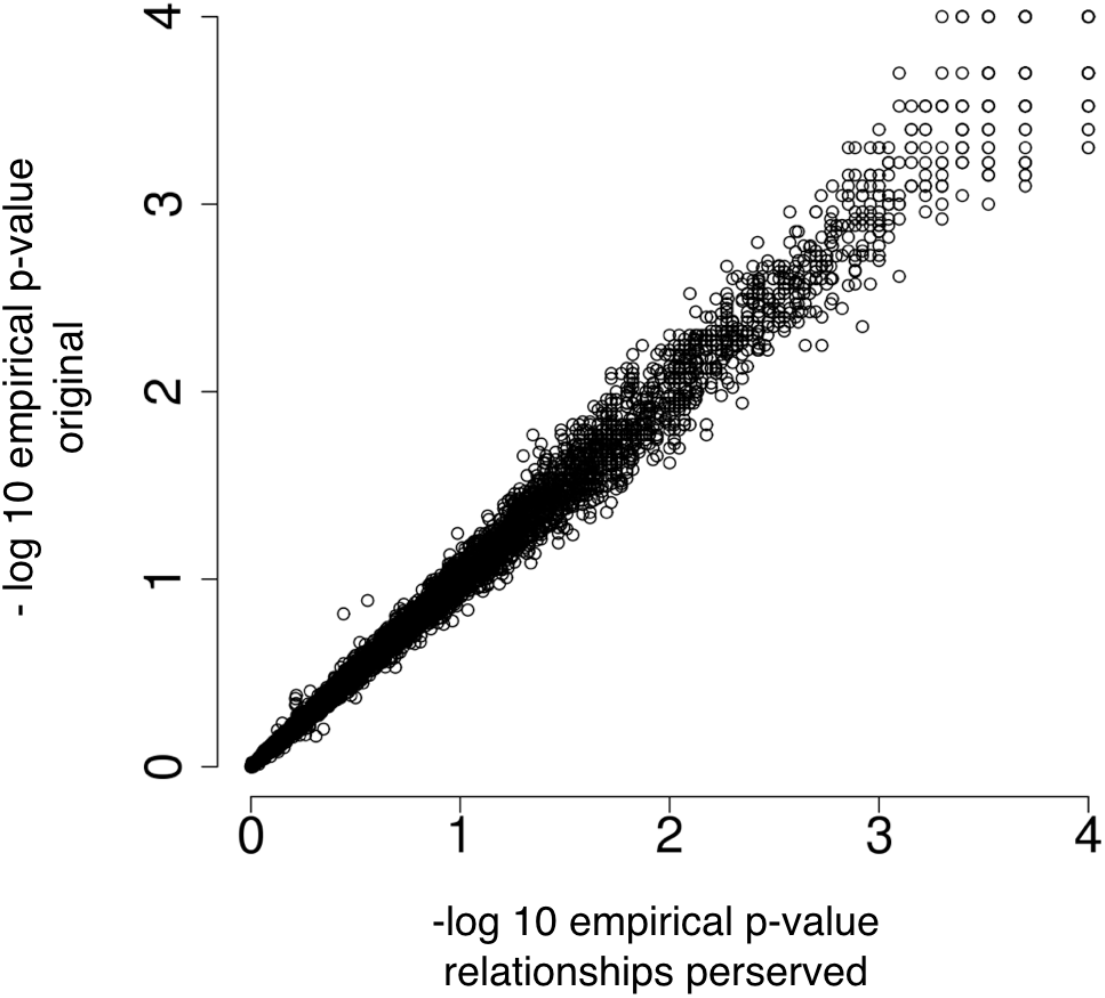
Comparison between transcript-relationship-preserving permutations with transcript-independent permutations. Comparison between empirical p-values between the approaches reveals similar values. The permutations that preserve the transcript-transcript relationship has 100 fewer significant genes at FDR < 0.05, but also had 234 fewer tested transcripts. Fewer transcripts were tested when performing permutations that swap the entire dataset (thereby preserving the transcript-transcript relationship) because there are genes that are missing in certain strains. Missing data is not handled by GEMMA, so these genes could not be tested. To avoid this, we performed permutations independently for each transcript in our analyses.

**Supplemental Figure 12:**
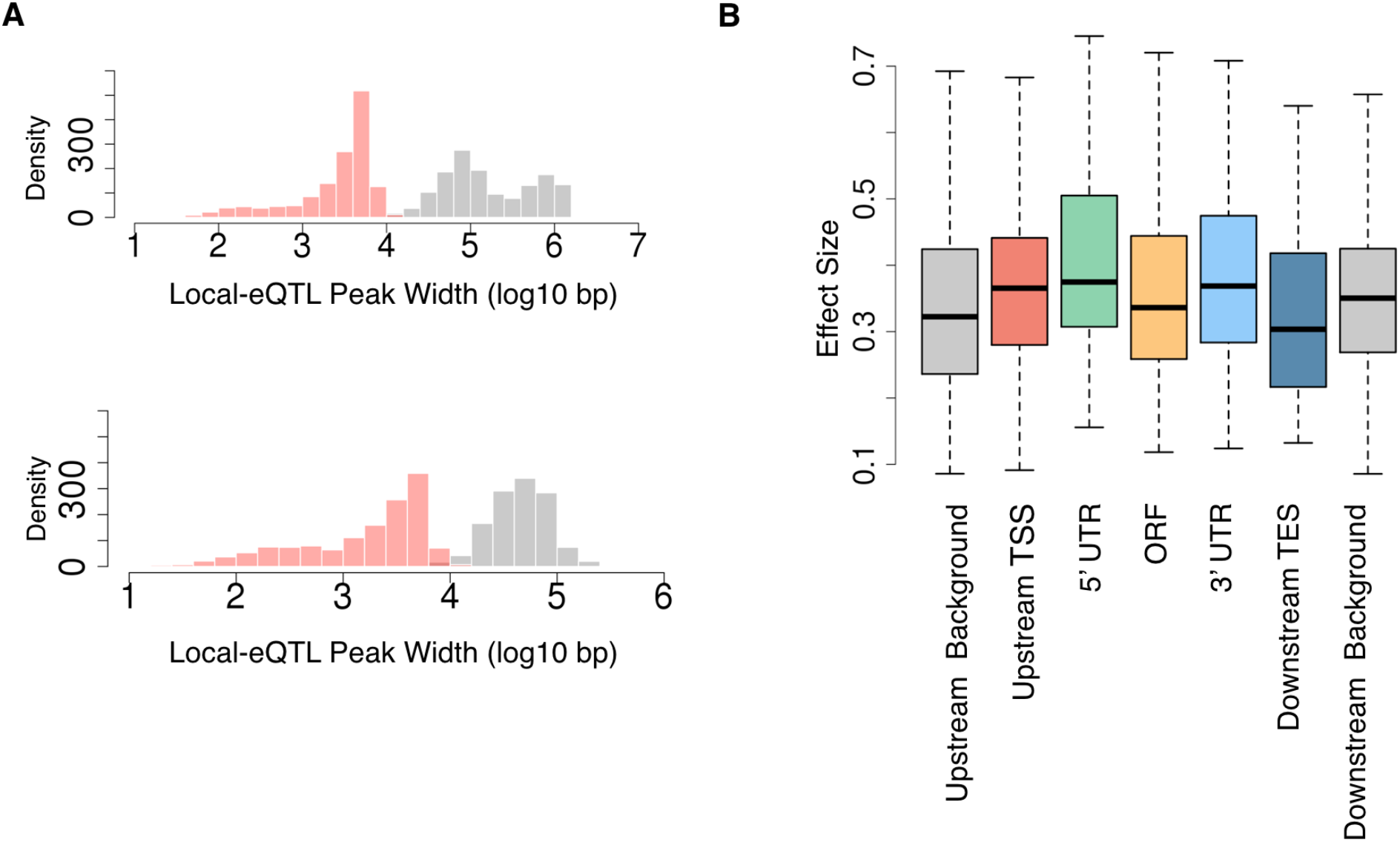
Local eQTL resolution and effect sizes of eQTLs according to location. A. The distribution of 2-LOD eQTL support interval from Ronald et al. (top, grey) and 1.5 LOD local eQTL support interval from Smith and Kruglyak 2008 (bottom, grey). Red distributions indicate the respective support interval observed in this study (2-LOD top and 1.5 LOD bottom). B. The distribution of effect sizes for eQTLs with each location type. The effect size is the absolute value of the beta value obtained from the GEMMA analysis.

**Supplemental Figure 13:**
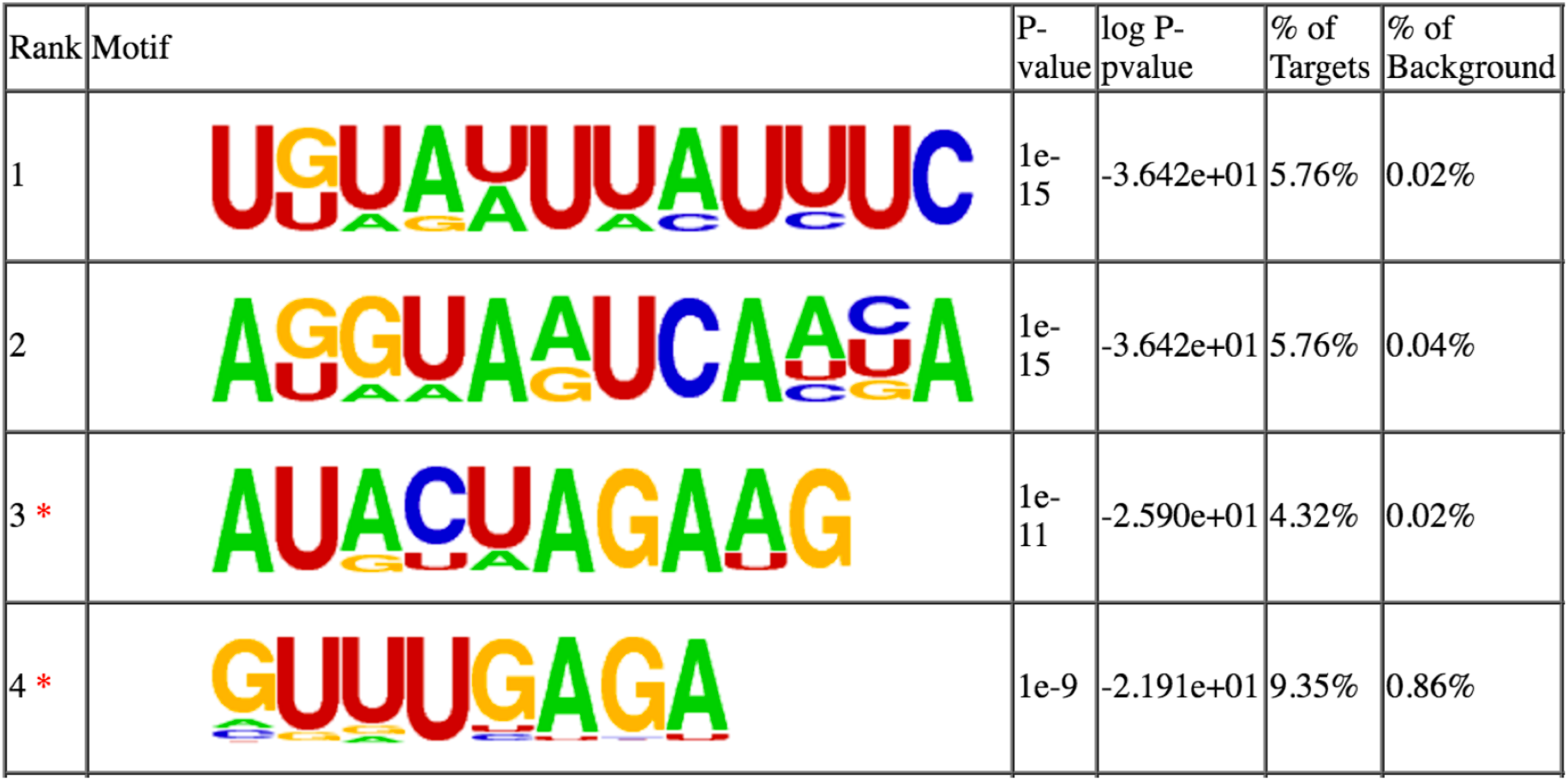
Homer enrichment result for the 3’ UTR eQTLs. The top four results from the HOMER output. The motifs without the “*” are significant after multiple hypothesis testing.

**Supplemental Figure 14:**
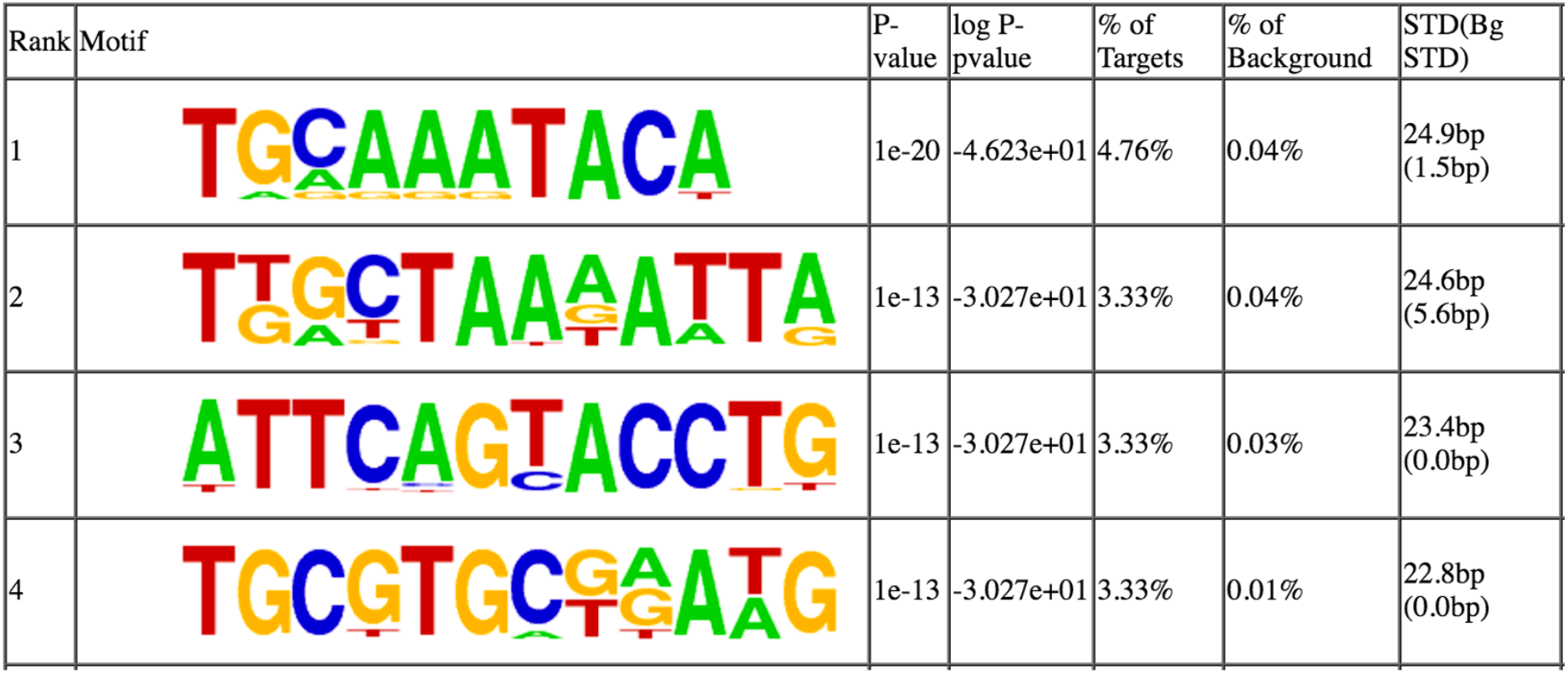
Homer enrichment result for the promoter regions. The top four results from the HOMER output. All four are significant after multiple hypothesis testing.

**Supplemental Figure 15:**
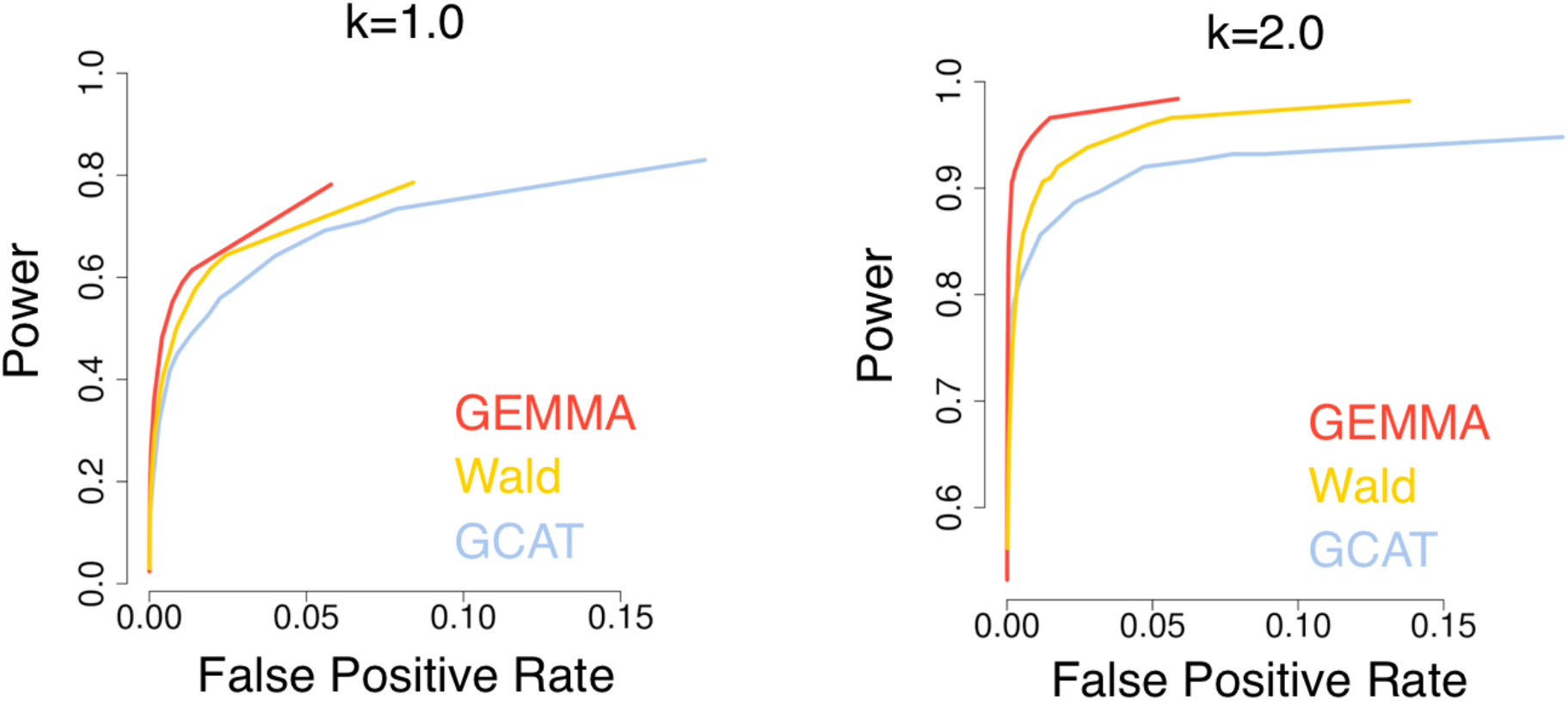
Power analysis of GWAS and population-structure correction. Receiver operating characteristic curve from simulations for the three methods that were tested.

**Supplemental Figure 16:**
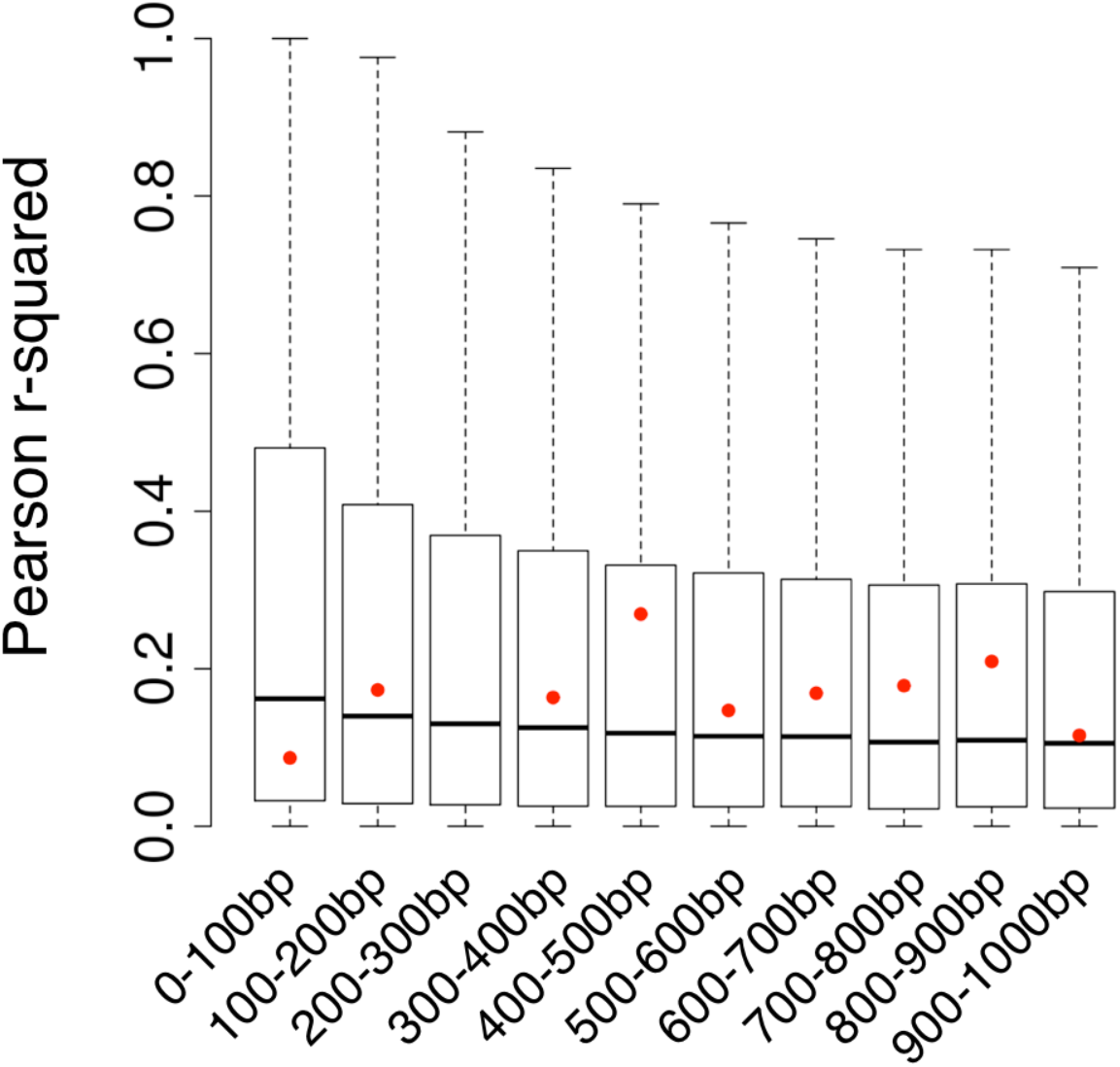
Assessment of genome-wide linkage disequilibrium. Genome-wide linkage disequilibrium was assessed by choosing 100,000 random SNPs (see Supplemental materials description). The distribution of the median linkage to the random SNPs within each window is shown by the boxplots. The median linkage to the chrXII: 830378 SNP is shown by the red dots. The missing value in the third window is because there were no SNPs within that window neighboring the SNP.

**Supplemental Figure 17:**
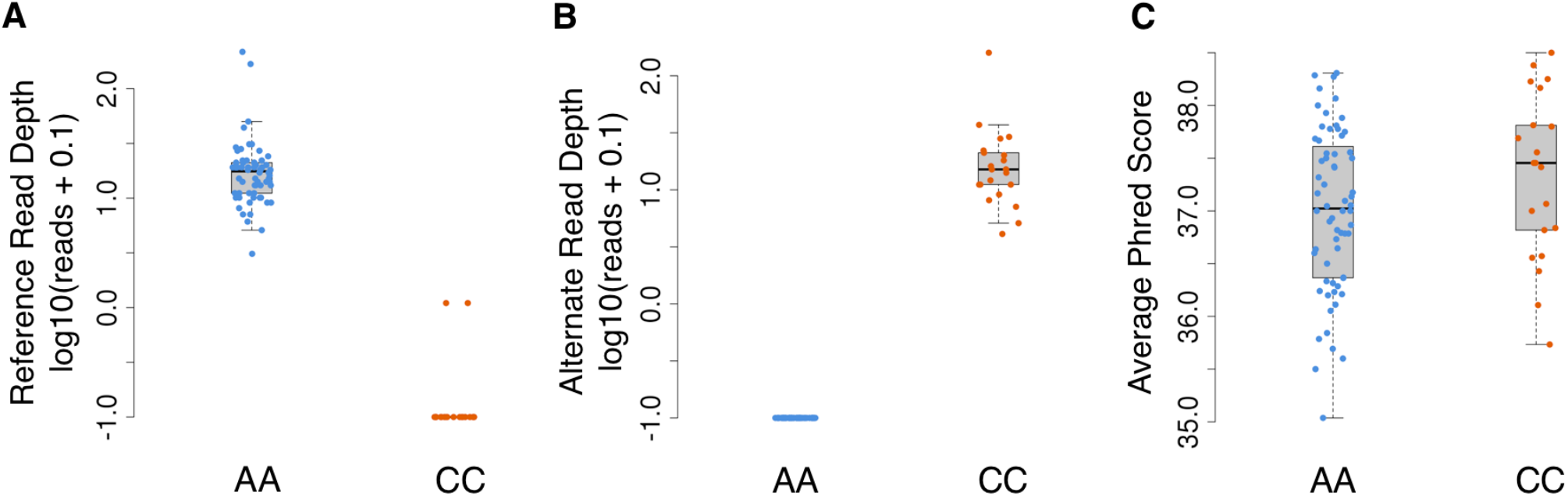
Assessment of the quality of the chrXI: 830378 SNP. (A) Number of reads with the reference allele for each strain, segregated by the final genotyping call for that strain. (B) Number of reads with the alternate allele for each strain, segregated by the final genotyping call for that strain. (C) Average phred score among the reads with the final-called allele for each strain.

**Supplemental Figure 18:**
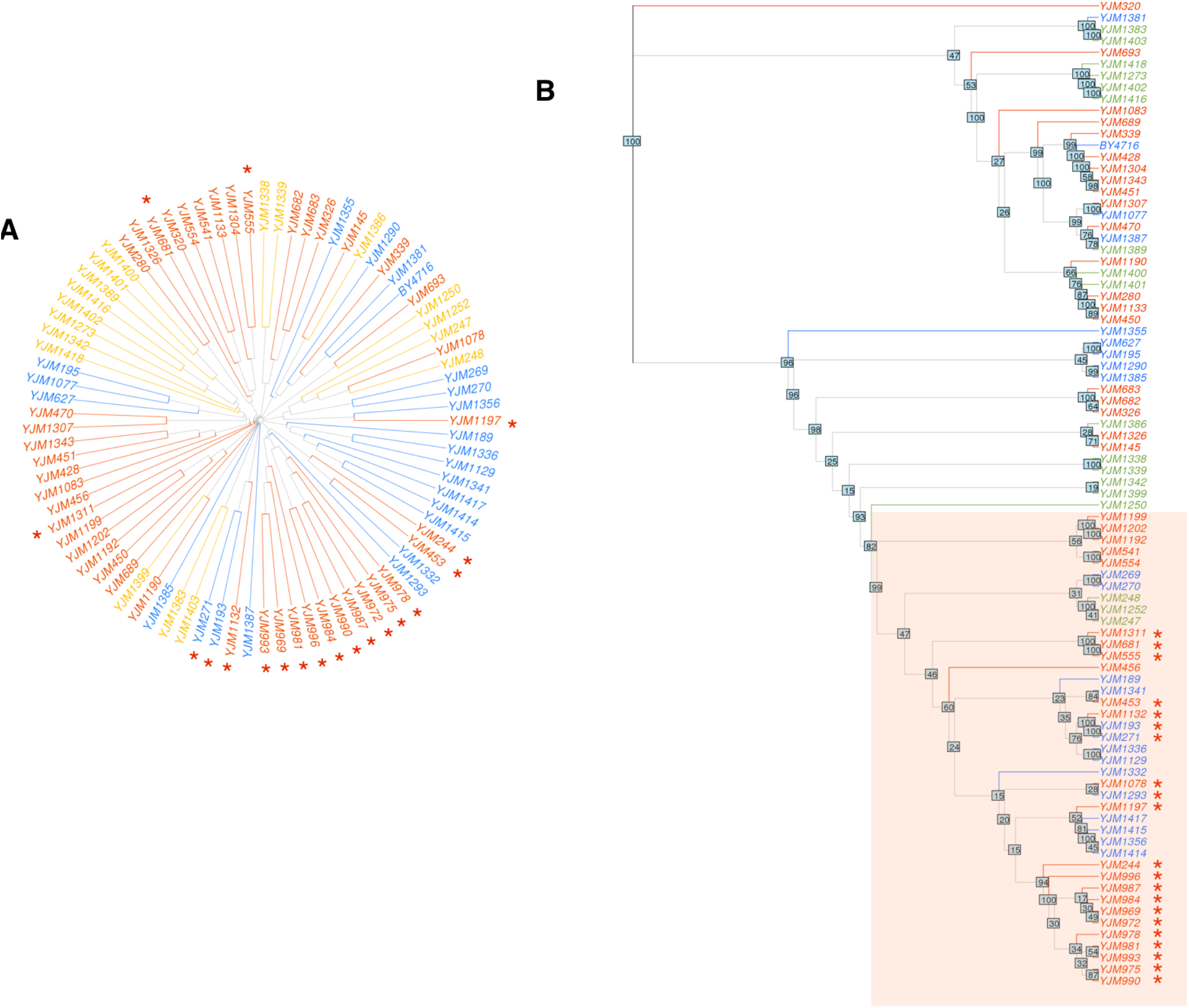
Regional neighbor-joining tree of region surrounding NIT3. (A) Phylogeny is from the genome-wide neighbor joining tree as shown in Figure 1. The red star indicates the strains with the clinical allele of the NIT3 upstream variant. (B) Neighbor-joining tree of the 50kb region surrounding NIT3. The consistency of the nodes across 100 rounds of bootstrapping is shown at each node. The red square indicates the smallest high-confidence group (>95 from bootstrapping) that contains all of the NIT3 clinical origin allele.

**Supplemental Figure 19:**
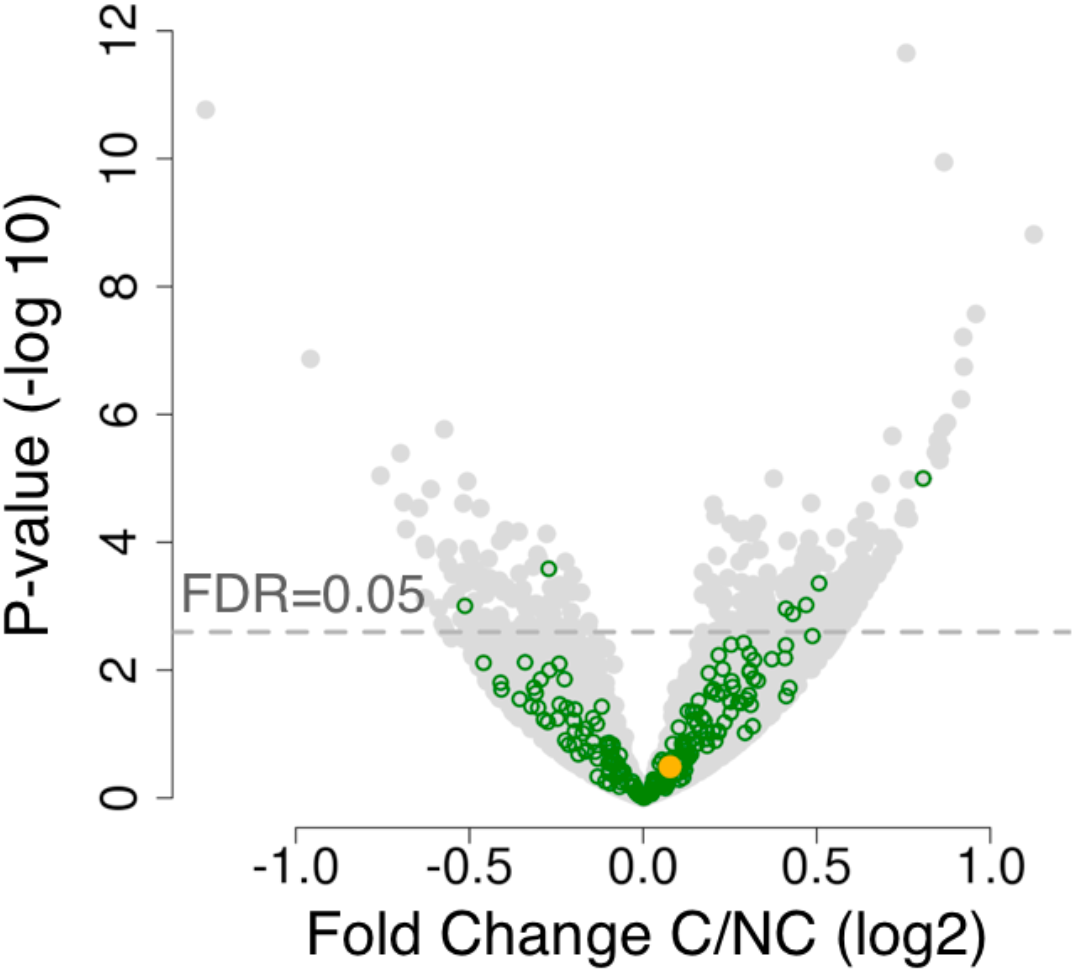
Differential expression between clinical and non-clinical strains. Volcano plot of differential expression between clinical and non-clinical strains. NIT3 is labeled in orange. FDR line is calculated from Benjamini-Hochberg. Green circles indicate biofilm suppressor genes.

## SUPPLEMENTAL TABLES

<u>Supplemental Table 1:</u> Description of the 85 isolates analyzed in this study.

<u>Supplemental Table 2: local-</u>eQTL results

<u>Supplemental Table 3:</u> Differential expression results between clinical and non-clinical strains.

## SUPPLEMENTARY TEXT

### Identifying absent genes

Strains of *S. cerevisiae* exhibit significant differences in the presence and absence of genes (Bergstrom et al. 2014; Strope et al. 2015). These effects can increase noise in eQTL mapping because the presence and absence of genes will affect the observed expression levels without the existence of a regulatory variant. Here we attempt to identify the absence of genes to aid our mapping of eQTLs. Uneven coverage in the DNA sequencing dataset precluded the estimation of copy number variation, however absence can be assessed because it will affect both the RNA and DNA in a predictable manner. For each strain, we first marked genes as potentially absent if the DNA-seq mapping had low coverage across that gene (RPKM < 1). To decrease false positives, we performed the same analysis on the Strope et al. sequencing data, and required genes to be called absent in both analyses. We then confirmed with the RNA-seq data to ensure that there was also no expression from these genes. Using the resulting list of absent genes, we confirmed two expected characteristics: 1. BY4716 is a strain that is nearly identical to the reference genome S288c, except for the designed deletion of one locus, *lys2* (Brachmann et al. 1998), which was the only gene we detected as absent. 2. We also observed that absent genes occurred primarily near chromosome ends, in agreement with previous observations (Brown et al. 2010; Bergstrom et al. 2014).

### eQTL replication with Skelly et al.

Concordance was assessed by first calculating variant-expression associations in Skelly et al. To calculate the variants, we identified all SNPs with MAF > 0.05. We then processed the RNA-seq data using the same approach as for the data generated in this study (quantile-normalization, fitting to standard normal, regressing out PEER factors), and tested for association with GEMMA.

To assess the agreement with our eQTLs, we compared the overall distribution of p-values and directionality. For each gene, we extracted the best variant-expression association found in our data. We then extracted the p-value and directionality of the same variant-expression pair in the Skelly et al. dataset. To compare the concordance in p-value, we set an empirical p-value cutoff to separate the gene-SNP pairs into two groups according to their empirical p-value in our data. We then compared the distribution of Skelly et al. p-values for each group using a Wilcoxon test, recording the p-value from this comparison. We then repeated this analysis using a range of empirical p-value cutoffs to separate the two groups (Supplemental Figure 10).

To compare the concordance in directionality, we sorted the gene-SNP pairs by their empirical p-value in this study. We then performed a sliding window analysis with a window size of 1000 genes. For each window, we used the binomial distribution to calculate the probability of observing at least as many gene-SNP pairs matching directionality in Skelly et al. and our dataset (compared to the 500 expected by chance). We then recorded the binomial test p-value and the median empirical p-value of the window (Supplemental Figure 10).

### Power analysis and population-structure correction

Using simulations, we compared the efficacy in correcting for population-structure between the Wald test, GEMMA, and GCAT (Zhou and Stephens 2012; Song et al. 2015). For each simulation, we selected a random SNP and simulated a phenotype with effect size k of 1.0 and 2.0 (equation for phenotype simulation according to (Long and Langley 1999)).The simulations were performed as described in (Strope et al. 2015) and (Connelly and Akey 2012). True positives were simulated by choosing a random variant that is local to the gene and simulating expression values for all strains to match the chosen effect size. False positives were simulated by choosing a random variant that is *not local* to the gene and simulating expression values with the chosen effect size. We performed simulations with various numbers of true positives and false positives and compared the false positive rate with the true positive rate for several alpha values.

### Linkage disequilibrium decay

A gradual decrease of significant p-values is commonly seen in GWAS analyses performed in humans and mice (Bennett et al. 2010; Civelek and Lusis 2014). However, we did not observe this pattern nearby the strongest associating SNP in this study. The lack of pattern is concordant with previous *S. cerevisiae* GWAS analyses using a tiling array (Muller et al. 2011). To further confirm that this lack of neighboring p-value decay is not specific to this SNP (which could indicate sequencing errors), we analyzed the linkage disequilibrium across all regions of the genome. We selected 10^4^ random SNPs across the genome, and calculated the median r^2^ (from the Pearson correlation) across 100 base-pair windows of increasing distance (Supplemental Figure 16). We then performed the same calculation for the SNP of interest (chrXII:830378). We observed that the SNP of interest exhibits a similar level of linkage disequilibrium as the rest of the genome.

### Phylogenomic analyses

The genomic tree was constructed from the set of highly conserved regions (total 218 kb) selected by Strope et al. for phylogenomic analyses. To construct strain-specific sequences in those regions, variants were inserted into the S288c reference sequence. Indels were excluded to avoid the possibility of improper weighting at gaps. Neighbor joining trees were then created with the ape package using the default parameters (distance matrix = K80) (Paradis et al. 2004). The expression tree was constructed using RPKM data filtered for genes with at least ten strains greater than 0.1 RPKM. The RPKM data were then quantile-normalized and each gene was standardized. The distance matrix was constructed using Euclidean distance. The genomic tree of the NIT3 region was constructed using all SNPs within 50 kb of the SNP of interest (chrXII: 830378). Confidence in the trees was assessed using bootstrapping with the boot.phylo method in ape with 100 permutations.

### NIT3 SNP population variation

We investigated the population variation of this SNP relative to the genome-wide population structure calculated above. The clinical-origin-associated allele is distributed across a diverse set of strains, with the greatest enrichment in one clade (Supplemental Figure 18A). However because genome-wide phylogenies show only an average of many complex admixtures, we then performed a phylogenetic analysis restricted to the 100 kb surrounding *NIT3,* to investigate the history of this specific region. Using the *NIT3*-region tree, we identified a high-confidence group that completely contained the strains with the clinical origin variant (Supplemental Figure 18B). This suggests that the variant likely originated from a single mutation, as opposed to multiple independent mutations, which would have resulted in the variant being present on disparate haplotypes.

### Selection analysis

Testing for overall selection on eQTLs was performed by two approaches: 1. Comparing allele frequencies of the eQTLs and 2. Analyzing the presence and absence of eQTLs across subsets of genes. For comparing allele frequencies, we repeated the eQTL mapping specifically for this analysis using the nonparametric Wilcoxon rank sum test (using variants with minimum minor allele frequency of 0.1). This resulted in analysis of 907 eQTLs at FDR < 0.05. We performed 1000 permutations where the gene expression was permuted between isolates. From each of these permutations, we recorded the variant/gene pairs that had p-values lower than the same cutoff used in the actual Wilcoxon eQTL mapping, and calculated the MAF distribution for 1000 groups of 907 randomized eQTLs. The eQTL mapping of the permutations was created using the same kinship matrix and parameters as described in the eQTL mapping methods, with the exception of using the Wilcoxon rank sum test instead of GEMMA. To test for presence and absence differences of eQTLs among specific subsets of genes, we corrected for confounding effects due to expression level by matching genes of similar expression. We matched genes with similar expression by sorting all genes by mean expression and binning into groups of 100 genes.

### Motif analysis

We searched for motifs across the eQTLs found within 3’ UTRs using HOMER (Heinz et al. 2010). To prevent enrichment of motifs that are found across all 3’ UTRs regardless of eQTL presence, we used as background the best associating variant found within 3’ UTR in genes that had an eQTL empirical p-value > 0.3. As the foreground, we used the best associating variant found within 3’ UTR in genes with an eQTL empirical p-value < 0.1. Parameters used were “sacCer3”, “homerUtr3”, “-size 100” “-rna”. For the promoter region, the “-rna” flag was not used.

## REFERENCES

Artieri CG, Fraser HB. 2014. Evolution at two levels of gene expression in yeast. Genome research 24: 411–421.

Albert FW, Kruglyak L. 2015. The role of regulatory variation in complex traits and disease. Nature Reviews Genetics 16: 197–212.

Altshuler D, Daly MJ, and Lander ES. 2008. Genetic Mapping in Human Disease. Science 322: 881–888.

Battle A, Mostafavi S, Zhu X, Potash JB, Weissman MM, McCormick C, Haudenschild CD, Beckman KB, Shi J, Mei R et al. 2014. Characterizing the genetic basis of transcriptome diversity through RNA-sequencing of 922 individuals. Genome Research 24: 14–24.

Benjamini Y, Hochberg Y. 1995. Controlling the False Discovery Rate: A Practical and Powerful Approach to Multiple Testing. Journal of the Royal Statistical Society Series B (Methodological) 57: 289–300.

Bergstrom A, Simpson JT, Salinas F, Barre B, Parts L, Zia A, Nguyen Ba AN, Moses AM, Louis EJ, Mustonen V et al. 2014. A high-definition view of functional genetic variation from natural yeast genomes. Molecular Biology and Evolution 31: 872–888.

Bojsen R, Regenberg B, Folkesson A. 2014. Saccharomyces cerevisiae biofilm tolerance towards systemic antifungals depends on growth phase. BMC Microbiology 14: 305.

Brem RB, Kruglyak L. 2005. The landscape of genetic complexity across 5,700 gene expression traits in yeast. Proceedings of the National Academy of Sciences of the United States of America 102: 1572–1577.

Brem RB, Storey JD, Whittle J, Kruglyak L. 2005. Genetic interactions between polymorphisms that affect gene expression in yeast. Nature 436: 701–703.

Brem RB, Yvert G, Clinton R, Kruglyak L. 2002. Genetic dissection of transcriptional regulation in budding yeast. Science 296: 752–755.

Chang CC, Chow CC, Tellier LC, Vattikuti S, Purcell SM, Lee JJ. 2015. Second-generation PLINK: rising to the challenge of larger and richer datasets. GigaScience 4: 7.

Chang J, Zhou Y, Hu X, Lam L, Henry C, Green EM, Kita R, Kobor MS, Fraser HB. 2013. The molecular mechanism of a cis-regulatory adaptation in yeast. PLoS Genetics 9: e1003813.

Cherry JM, Hong EL, Amundsen C, Balakrishnan R, Binkley G, Chan ET, Christie KR, Costanzo MC, Dwight SS, Engel SR et al. 2012. Saccharomyces Genome Database: the genomics resource of budding yeast. Nucleic Acids Research 40: D700–705.

Cingolani P, Platts A, Wang le L, Coon M, Nguyen T, Wang L, Land SJ, Lu X, Ruden DM. 2012. A program for annotating and predicting the effects of single nucleotide polymorphisms, SnpEff: SNPs in the genome of *Drosophila melanogaster* strain w1118; iso-2; iso-3. Fly 6: 80–92.

Claussnitzer M, Dankel SN, Kim KH, Quon G, Meuleman W, Haugen C, Glunk V, Sousa IS, Beaudry JL, Puviindran V et al. 2015. FTO Obesity Variant Circuitry and Adipocyte Browning in Humans. The New England Journal of Medicine 373: 895–907.

Connallon T, Knowles LL. 2007. Recombination rate and protein evolution in yeast. BMC Evol Biol 7: 235.

Connelly CF, Akey JM. 2012. On the prospects of whole-genome association mapping in *Saccharomyces cerevisiae*. Genetics 191: 1345–1353.

Cox MP, Peterson DA, Biggs PJ. 2010. SolexaQA: At-a-glance quality assessment of Illumina second-generation sequencing data. BMC Bioinformatics 11: 485.

Denver DR, Morris K, Streelman JT, Kim SK, Lynch M, Thomas WK. 2005. The transcriptional consequences of mutation and natural selection in *Caenorhabditis elegans*. Nature Genetics 37: 544–548.

Dobin A, Davis CA, Schlesinger F, Drenkow J, Zaleski C, Jha S, Batut P, Chaisson M, Gingeras TR. 2013. STAR: ultrafast universal RNA-seq aligner. Bioinformatics 29: 15–21.

Dunn B, Richter C, Kvitek DJ, Pugh T, Sherlock G. 2012. Analysis of the *Saccharomyces cerevisiae* pangenome reveals a pool of copy number variants distributed in diverse yeast strains from differing industrial environments. Genome Research 22: 908–924.

Enache-Angoulvant A, Hennequin C. 2005. Invasive *Saccharomyces* infection: a comprehensive review. Clinical Infectious Diseases 41: 1559–1568.

Enard D, Messer PW, Petrov DA. 2014. Genome-wide signals of positive selection in human evolution. Genome Research 24: 885–895.

Fanning S, Mitchell AP. 2012. Fungal biofilms. PLoS Pathogens 8: e1002585.

Flint J and Eskin E. 2012. Genome-wide association studies in mice. Nature Reviews Genetics 13: 807–817.

Fraser HB. 2011. Genome-wide approaches to the study of adaptive gene expression evolution. BioEssays 33: 469–477.

Fraser HB, 2013. Gene expression drives local adaptation in humans. Genome Research 23: 1089–1096.

Fraser HB, Babak T, Tsang J, Zhou Y, Zhang B, Mehrabian M, Schadt EE. 2011. Systematic detection of polygenic cis-regulatory evolution. PLoS Genet. 7: e1002023.

Fraser HB, Hirsh AE, Giaever G, Kumm J, Eisen MB. 2004. Noise minimization in eukaryotic gene expression. PLoS Biology 2: e137.

Fraser HB, Levy S, Chavan A, Shah HB, Perez JC, Zhou Y, Siegal ML, Sinha H. 2012. Polygenic cis-regulatory adaptation in the evolution of yeast pathogenicity. Genome Research 22: 1930–1939.

Fraser HB, Moses AM, Schadt EE. 2010. Evidence for widespread adaptive evolution of gene expression in budding yeast. Proceedings of the National Academy of Sciences of the United States of America 107: 2977–2982.

Freeberg MA, Han T, Moresco JJ, Kong A, Yang YC, Lu ZJ, Yates JR, Kim JK. 2013. Pervasive and dynamic protein binding sites of the mRNA transcriptome in *Saccharomyces cerevisiae*. Genome Biology 14: R13.

Garrison E, Marth G. 2012. Haplotype-based variant detection from short-read sequencing. ArXiv e-prints 1207.

Gerber AP, Herschlag D, Brown PO. 2004. Extensive association of functionally and cytotopically related mRNAs with Puf family RNA-binding proteins in yeast. PLoS Biology 2: E79.

Gusev A, Ko A, Shi H, Bhatia G, Chung W, Penninx BW, Jansen R, de Geus EJ, Boomsma DI, Wright FA et al. 2016. Integrative approaches for large-scale transcriptome-wide association studies. Nature Genetics 48: 245–252.

Hogan GJ, Brown PO, Herschlag D. 2015. Evolutionary Conservation and Diversification of Puf RNA Binding Proteins and Their mRNA Targets. PLoS Biology 13: e1002307.

Huang W, Carbone MA, Magwire MM, Peiffer JA, Lyman RF, Stone EA, Anholt RR, Mackay TF. 2015. Genetic basis of transcriptome diversity in Drosophila melanogaster. Proceedings of the National Academy of Sciences of the United States of America 112: E6010–6019.

Johnson WE, Li C, and Rabinovic A. 2007. Adjusting batch effects in microarray expression data using empirical Bayes methods. Biostatistics 8: 118–127.

Josephs EB, Lee YW, Stinchcombe JR, Wright SI. 2015. Association mapping reveals the role of purifying selection in the maintenance of genomic variation in gene expression. Proceedings of the National Academy of Sciences of the United States of America 112: 15390–15395.

Leek JT and Storey JD. 2007. Capturing Heterogeneity in Gene Expression Studies by Surrogate Variable Analysis. Plos Genetics 3: e161.

Li H, Handsaker B, Wysoker A, Fennell T, Ruan J, Homer N, Marth G, Abecasis G, Durbin R, Genome Project Data Processing S. 2009. The Sequence Alignment/Map format and SAMtools. Bioinformatics 25: 2078–2079.

Li Y, Willer CJ, Ding J, Scheet P, Abecasis GR. 2010. MaCH: using sequence and genotype data to estimate haplotypes and unobserved genotypes. Genetic Epidemiology 34: 816–834.

Liti G, Carter DM, Moses AM, Warringer J, Parts L, James SA, Davey RP, Roberts IN, Burt A, Koufopanou V et al. 2009. Population genomics of domestic and wild yeasts. Nature 458: 337–341.

Love MI, Huber W, Anders S. 2014. Moderated estimation of fold change and dispersion for RNA-seq data with DESeq2. Genome Biology 15: 550.

Lunter G, Goodson M. 2011. Stampy: a statistical algorithm for sensitive and fast mapping of Illumina sequence reads. Genome Research 21: 936–939.

Martin HC, Roop JI, Schraiber JG, Hsu TY, Brem RB. 2012. Evolution of a membrane protein regulon in Saccharomyces. Molecular Biology and Evolution 29: 1747–1756.

McManus CJ, May GE, Spealman P, Shteyman A. 2014. Ribosome profiling reveals post-transcriptional buffering of divergent gene expression in yeast. Genome research 24: 422–430.

Muller LA, Lucas JE, Georgianna DR, McCusker JH. 2011. Genome-wide association analysis of clinical vs. nonclinical origin provides insights into *Saccharomyces cerevisiae* pathogenesis. Molecular Ecology 20: 4085–4097.

Munoz P, Bouza E, Cuenca-Estrella M, Eiros JM, Perez MJ, Sanchez-Somolinos M, Rincon C, Hortal J, Pelaez T. 2005. Saccharomyces cerevisiae fungemia: an emerging infectious disease. Clinical Infectious Diseases 40: 1625–1634.

Musunuru K, Strong A, Frank-Kamenetsky M, Lee NE, Ahfeldt T, Sachs KV, Li X, Li H, Kuperwasser N, Ruda VM et al. 2010. From noncoding variant to phenotype via *SORT1* at the 1p13 cholesterol locus. Nature 466: 714–719.

Naranjo S, Smith JD, Artieri CG, Zhang M, Zhou Y, Palmer ME, Fraser HB. 2015. Dissecting the Genetic Basis of a Complex cis-Regulatory Adaptation. PLoS Genetics 11: e1005751.

Nobile CJ, Fox EP, Nett JE, Sorrells TR, Mitrovich QM, Hernday AD, Tuch BB, Andes DR, Johnson AD. 2012. A recently evolved transcriptional network controls biofilm development in *Candida albicans*. Cell 148: 126–138.

Nobile CJ, Johnson AD. 2015. Candida albicans Biofilms and Human Disease. Annual Review of Microbiology 69: 71–92.

Pace HC, Brenner C. 2001. The nitrilase superfamily: classification, structure and function. Genome Biology 2: REVIEWS0001.

Pai AA, Cain CE, Mizrahi-Man O, De Leon S, Lewellen N, Veyrieras JB, Degner JF, Gaffney DJ, Pickrell JK, Stephens M et al. 2012. The contribution of RNA decay quantitative trait loci to inter-individual variation in steady-state gene expression levels. PLoS Genetics 8: e1003000.

Pelechano V, Wei W, Steinmetz LM. 2013. Extensive transcriptional heterogeneity revealed by isoform profiling. Nature 497: 127–131.

Purcell S, Neale B, Todd-Brown K, Thomas L, Ferreira MA, Bender D, Maller J, Sklar P, de Bakker PI, Daly MJ et al. 2007. PLINK: a tool set for whole-genome association and population-based linkage analyses. American Journal of Human Genetics 81: 559–575.

Reynolds TB, Fink GR. 2001. Bakers’ yeast, a model for fungal biofilm formation. Science 291: 878–881.

Rhee HS, Pugh BF. 2012. Genome-wide structure and organization of eukaryotic pre-initiation complexes. Nature 483: 295–301.

Rifkin SA, Houle D, Kim J, White KP. 2005. A mutation accumulation assay reveals a broad capacity for rapid evolution of gene expression. Nature 438: 220–223.

Rockman MV, Kruglyak L. 2006. Genetics of global gene expression. Nature Reviews Genetics 7: 862–872.

Ronald J, Akey JM. 2007. The evolution of gene expression QTL in Saccharomyces cerevisiae. PloS One 2: e678.

Ronald J, Brem RB. Whittle J, Kruglyak L. 2005. Local regulatory variation in Saccharomyces cerevisiae. PLoS Genetics 1: e25.

Roop JI, Chang KC, Brem RB, 2016. Polygenic evolution of a sugar specialization trade-off in yeast. Nature 530: 336–339.

Schacherer J, Shapiro JA, Ruderfer DM, Kruglyak L. 2009. Comprehensive polymorphism survey elucidates population structure of Saccharomyces cerevisiae. Nature 458: 342–345.

Schaub MA, Boyle AP, Kundaje A, Batzoglou S, Snyder M. 2012. Linking disease associations with regulatory information in the human genome. Genome Research 22: 1748–1759.

Scherz K, Andersen, Bojsen R, Gro L, Rejkjaer, Sorensen, Weiss M, Nielsen, Lisby M, Folkesson A et al. 2014. Genetic basis for Saccharomyces cerevisiae biofilm in liquid medium. G3 4: 1671–1680.

Sellis D, Kvitek DJ, Dunn B, Sherlock G, Petrov DA. 2016. Heterozygote Advantage Is a Common Outcome of Adaptation in Saccharomyces cerevisiae. Genetics 203: 1401–1413.

Sharon E, Kalma Y, Sharp A, Raveh-Sadka T, Levo M, Zeevi D, Keren L, Yakhini Z, Weinberger A, Segal E. 2012. Inferring gene regulatory logic from high-throughput measurements of thousands of systematically designed promoters. Nature Biotechnology 30: 521–530.

Skelly DA, Merrihew GE, Riffle M, Connelly CF, Kerr EO, Johansson M, Jaschob D, Graczyk B, Shulman NJ, Wakefield J et al. 2013. Integrative phenomics reveals insight into the structure of phenotypic diversity in budding yeast. Genome Research 23: 1496–1504.

Smith EN, Kruglyak L. 2008. Gene-environment interaction in yeast gene expression. PLoS Biology 6: e83.

Stegle O, Parts L, Durbin R, Winn J. 2010. A Bayesian framework to account for complex non-genetic factors in gene expression levels greatly increases power in eQTL studies. PLoS Computational Biology 6: e1000770.

Stranger BE, Montgomery SB, Dimas AS, Parts L, Stegle O, Ingle CE, Sekowska M, Smith GD, Evans D, Gutierrez-Arcelus M et al. 2012. Patterns of cis regulatory variation in diverse human populations. PLoS Genetics 8: e1002639.

Strope PK, Skelly DA, Kozmin SG, Mahadevan G, Stone EA, Magwene PM, Dietrich FS, McCusker JH. 2015.The 100-genomes strains, an *S. cerevisiae* resource that illuminates its natural phenotypic and genotypic variation and emergence as an opportunistic pathogen. Genome Research 25: 762–774.

Tirosh I, Barkai N. 2008. Evolution of gene sequence and gene expression are not correlated in yeast. Trends in Genetics 24: 109–113.

Tirosh I, Reikhav S, Levy AA, Barkai N. 2009. A yeast hybrid provides insight into the evolution of gene expression regulation. Science 324: 659–662.

Tung J, Zhou X, Alberts SC, Stephens M, Gilad Y. 2015. The genetic architecture of gene expression levels in wild baboons. eLife 4: e04729.

Uppuluri P, Chaturvedi AK, Srinivasan A, Banerjee M, Ramasubramaniam AK, Kohler JR, Kadosh D, Lopez-Ribot JL. 2010. Dispersion as an important step in the Candida albicans biofilm developmental cycle. PLoS Pathogens 6: e1000828.

Uppuluri P, Lopez-Ribot JL. 2016. Go Forth and Colonize: Dispersal from Clinically Important Microbial Biofilms. PLoS Pathogens 12: e1005397.

Vandenbosch D, De Canck E, Dhondt I, Rigole P, Nelis HJ, Coenye T. 2013. Genomewide screening for genes involved in biofilm formation and miconazole susceptibility in Saccharomyces cerevisiae. FEMS Yeast Research 13: 720–730.

Veyrieras JB, Gaffney DJ, Pickrell JK. Gilad Y, Stephens M, Pritchard JK. 2012. Exon-specific QTLs skew the inferred distribution of expression QTLs detected using gene expression array data. PloS One 7: e30629.

Veyrieras JB, Kudaravalli S, Kim SY, Dermitzakis ET, Gilad Y, Stephens M, Pritchard JK, 2008. High-resolution mapping of expression-QTLs yields insight into human gene regulation. PLoS Genetics 4:e1000214.

Xu W, Solis NV, Ehrlich RL, Woolford CA, Filler SG, and Mitchell AP. 2015. Activation and Alliance of Regulatory Pathways in *C. albicans* during Mammalian Infection. Plos Biology 13: e1002076.

Zhou X, Stephens M. 2012. Genome-wide efficient mixed-model analysis for association studies. Nature Genetics 44: 821–824.

Zhu Z, Zhang F, Hu H, Bakshi A, Robinson MR, Powell JE, Montgomery GW, Goddard ME, Wray NR, Visscher PM et al. 2016. Integration of summary data from GWAS and eQTL studies predicts complex trait gene targets. Nature Genetics 48: 481–487.

## SUPPLEMENTAL MATERIAL CITATIONS

Bennett BJ, Farber CR, Orozco L, Kang HM, Ghazalpour A, Siemers N, Neubauer M, Neuhaus I, Yordanova R, Guan B et al. 2010. A high-resolution association mapping panel for the dissection of complex traits in mice. Genome Research 20: 281–290.

Brachmann CB, Davies A, Cost GJ, Caputo E, Li J, Hieter P, Boeke JD. 1998. Designer deletion strains derived from Saccharomyces cerevisiae S288C: a useful set of strains and plasmids for PCR-mediated gene disruption and other applications. Yeast 14: 115–132.

Brown CA, Murray AW, Verstrepen KJ. 2010. Rapid expansion and functional divergence of subtelomeric gene families in yeasts. Current Biology: CB 20: 895–903.

Civelek M, Lusis AJ. 2014. Systems genetics approaches to understand complex traits. Nature Reviews Genetics 15: 34–48.

Connelly CF, Akey JM. 2012. On the prospects of whole-genome association mapping in Saccharomyces cerevisiae. Genetics 191: 1345–1353.

Heinz S, Benner C, Spann N, Bertolino E, Lin YC, Laslo P, Cheng JX, Murre C, Singh H, Glass CK. 2010. Simple combinations of lineage-determining transcription factors prime cis-regulatory elements required for macrophage and B cell identities. Molecular Cell 38: 576–589.

Long AD, Langley CH. 1999. The power of association studies to detect the contribution of candidate genetic loci to variation in complex traits. Genome Research 9: 720–731.

Muller LA, Lucas JE, Georgianna DR, McCusker JH. 2011. Genome-wide association analysis of clinical vs. nonclinical origin provides insights into Saccharomyces cerevisiae pathogenesis. Molecular Ecology 20: 4085–4097.

Paradis E, Claude J, Strimmer K. 2004. APE: Analyses of Phylogenetics and Evolution in R language. Bioinformatics 20: 289–290.

Song M, Hao W, Storey JD. 2015. Testing for genetic associations in arbitrarily structured populations. Nature Genetics 47: 550–554.

Strope PK, Skelly DA, Kozmin SG, Mahadevan G, Stone EA, Magwene PM, Dietrich FS, McCusker JH. 2015. The 100-genomes strains, an S. cerevisiae resource that illuminates its natural phenotypic and genotypic variation and emergence as an opportunistic pathogen. Genome Research 25: 762–774.

Zhou X, Stephens M. 2012. Genome-wide efficient mixed-model analysis for association studies. Nature genetics 44: 821–824.

